# miR-29a-3p, a new myokine orchestrating resistance exercise via coordinated metabolic responses

**DOI:** 10.1101/2024.05.14.592416

**Authors:** Paola Pinto-Hernandez, Manuel Fernandez-Sanjurjo, Daan Paget, Xurde M Caravia, David Roiz-Valle, Juan Castilla-Silgado, Sergio Diez-Robles, Almudena Coto-Vilcapoma, Pau Gama, Pablo M Garcia-Roves, Carlos Lopez-Otin, Juleen Zierath, Anna Krook, Benjamin Fernandez-Garcia, Cristina Tomas-Zapico, Eduardo Iglesias-Gutierrez

## Abstract

It remains unclear whether the adaptive response to different exercise models is mediated by EV miRNAs released from skeletal muscle and their functional metabolic role. We sequenced miRNA-loaded plasma EVs obtained from resting mice after 4-weeks endurance or resistance training. Resistance exercise increased the expression of a 11-miRNA profile grouped into two functional clusters. Using both genetically modified animal models and in vitro approaches, we have identified miR-29a-3p as a novel myokine secreted into the bloodstream as EV cargo by contracting skeletal muscle. It is a cornerstone in the adaptation to resistance training by mediating the coordinated expression and secretion of other miRNAs and affecting muscle mass development and energy metabolism in muscle and liver. Taken together, our study suggests a coordinating and determinant role of miR-29a-3p in the response and adaptation to resistance training, possibly due to its role as a myokine through its regulatory role in energy metabolism.

## INTRODUCTION

The benefits of regular exercise for improving quality of life, as well as for the prevention and/or adjuvant treatment of chronic diseases, have been widely demonstrated ^1^. The systemic response to exercise involves not only skeletal muscle, as the primary responder to this stimulus, but also different tissues, including liver, heart, adipose tissue, kidneys, skin, pancreas, or brain ^2^. However, the molecular adaptations underlying this finely orchestrated multisystemic response, mediated by an intense inter-tissue crosstalk, have not been fully explored.

Several authors have reported the release of extracellular vesicles (EVs) into the systemic circulation from different tissues and blood cell types, that may participate in cell communication signaling during the response and adaptation to exercise ^3^. Thus, in 2018, Whitham et al. ^4^ described intercellular interactions between skeletal muscle and liver, mediated by protein transport via EV trafficking, in response to an acute bout of endurance exercise. Interestingly, EVs internalization by recipient cells implies the ability of their cargo to modify and/or induce different physiological pathways ^5,6^. As reviewed by Estébanez et al. ^7^ and Darragh et al. ^8^, changes not only in EV abundance but also in cargo profile has been described in response to acute exercise of different modalities and at rest in trained individuals, both in rodents and humans. Apart from proteins, the EV cargo comprises a variety of other macromolecules, including microRNAs (miRNAs) ^9^. MiRNAs are small non-coding RNA molecules (≈22 nucleotides) with a post-transcriptional gene expression regulatory function, by promoting mRNA degradation or by repressing protein translation ^10^. Therefore, miRNAs contained in EVs may modulate the gene expression and phenotype of distant recipient cells ^11^. Skeletal muscle acts as an important endocrine organ and it may also has the capacity to release miRNA-loaded EVs into the systemic circulation during exercise ^12^, while the liver has been described as the main receptor of plasma EVs ^4^. Therefore, it is worth considering whether the adaptive response to different models of regular exercise is, at least in part, mediated by EV miRNAs released by skeletal muscle and its functional metabolic interaction with liver.

In this study, we report the analysis of miRNA-loaded plasma EVs obtained from mice in resting state after 4-weeks of endurance or resistance training. We found that, although both exercise modalities show a common EV miRNA signature, resistance exercise exerts a higher and significant increase in this profile. Using in silico approaches, we determined that these EV miRNAs are mainly organized in two functional clusters. Based on these data and using both genetically modified animal models and in vitro approaches, we have identified miR-29a-3p as a new myokine secreted to bloodstream as EV cargo by contracting skeletal muscle playing a role as a cornerstone in the adaptation to resistance training through its regulatory role in muscle mass development and in energy metabolism pathways, both in muscle and liver.

## RESULTS

### Next-Generation Sequencing revealed a specific EV-miRNA profile in response to training functionally grouped in two clusters

With the aim of screening the resting profile of miRNAs contained in plasma EVs in the trained state, an exercise intervention was designed and implemented in C57BL6N 12-week-old male mice (n=26) (Figure 1A). The intervention consisted of 4 weeks of endurance (END) or resistance (RES) training, following previously used protocols ^13,14^. Another group of animals did not undergo any training during this period (CON). Endurance and resistance maximal capacities were determined by incremental tests (see STAR Methods), before and after the intervention. END and RES animals improved the capacity trained, while CON animals maintained their resistance capacity but worsened their aerobic endurance capacity (Figures 1B-D). Blood and tissue samples were collected 24 hours after the last training session. Plasma was immediately separated by centrifugation, aliquoted, and frozen.

**Figure 1.**
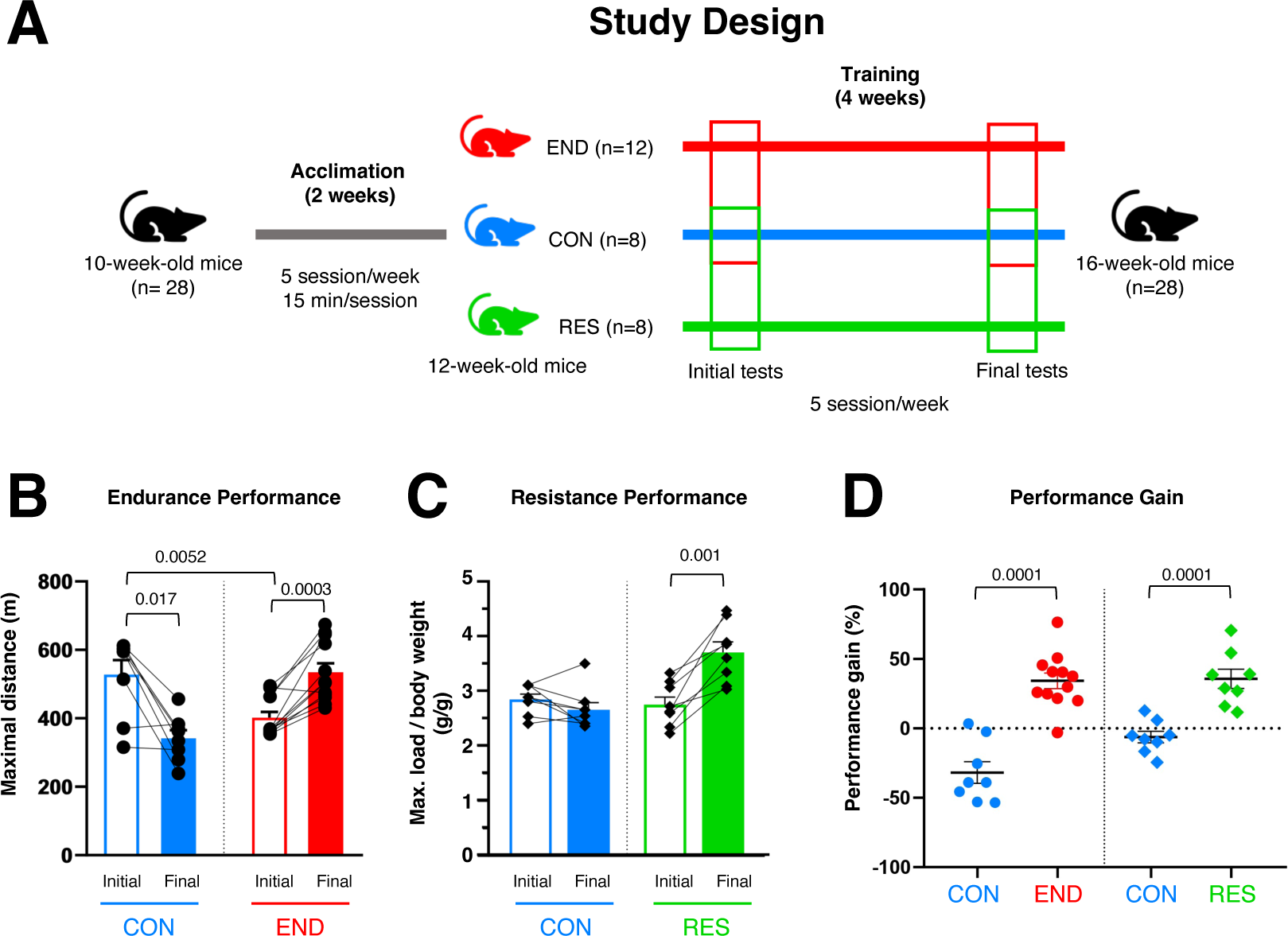
Endurance and resistance performance in WT mice. (A) Schematic representation of the exercise intervention (sedentary, CON; endurance, END; and resistance, RES). (B) Initial and final endurance performance of CON and END. (C) Initial and final resistance performance of CON and RES. (D) Performance gain (%) of CON (n=8), END (n=12), and RES (n=8), measured as percent variation. Data are presented as mean ± SEM. Each dot represents one mouse. Statistical significance measured by Student’s t-test for paired samples and for independent samples.

Since miRNAs present in EVs have been shown to participate in systemic crosstalk during exercise ^2,15^, total RNA, including small RNAs, was extracted from previously isolated EVs of pooled plasma samples (*n*=5/group) (Figure S1). Short RNA next generation sequencing (NGS) was then performed. We identified 223 miRNAs with a minimum expression of 1 Transcripts Per Kilobase Million (TPM) in at least one sample (http://doi.org/10.5281/zenodo.11184909). From them, 19 miRNAs with at least 3500 Reads Per Million (RPM) and a log_2_ fold change (FC) of ± 0.5 versus CON pool were validated by RT-qPCR. This internal validation was performed individually on the same samples per group used for NGS analysis.

A specific signature of 11 training-associated miRNAs was described as upregulated in END and RES mice regarding CON (Figure 2A). Ten of them were significantly increased in RES regarding CON mice: mmu-miR-10b-5p, mmu-miR-30a-5p, mmu-miR-126a-3p, mmu-miR-126a-5p, mmu-let-7a-5p, mmu-miR-29a-3p, mmu-miR-143-3p, mmu-let-7i-5p, mmu-miR-142a-5p and mmu-let-7c-5p, whereas mmu-let-7f-5p showed a notable, although not significant, increase (p=0.056) (Figure 2B). Increased levels of these miRNAs were also observed in END, although with no significant differences from CON (Figure 2B).

**Figure 2.**
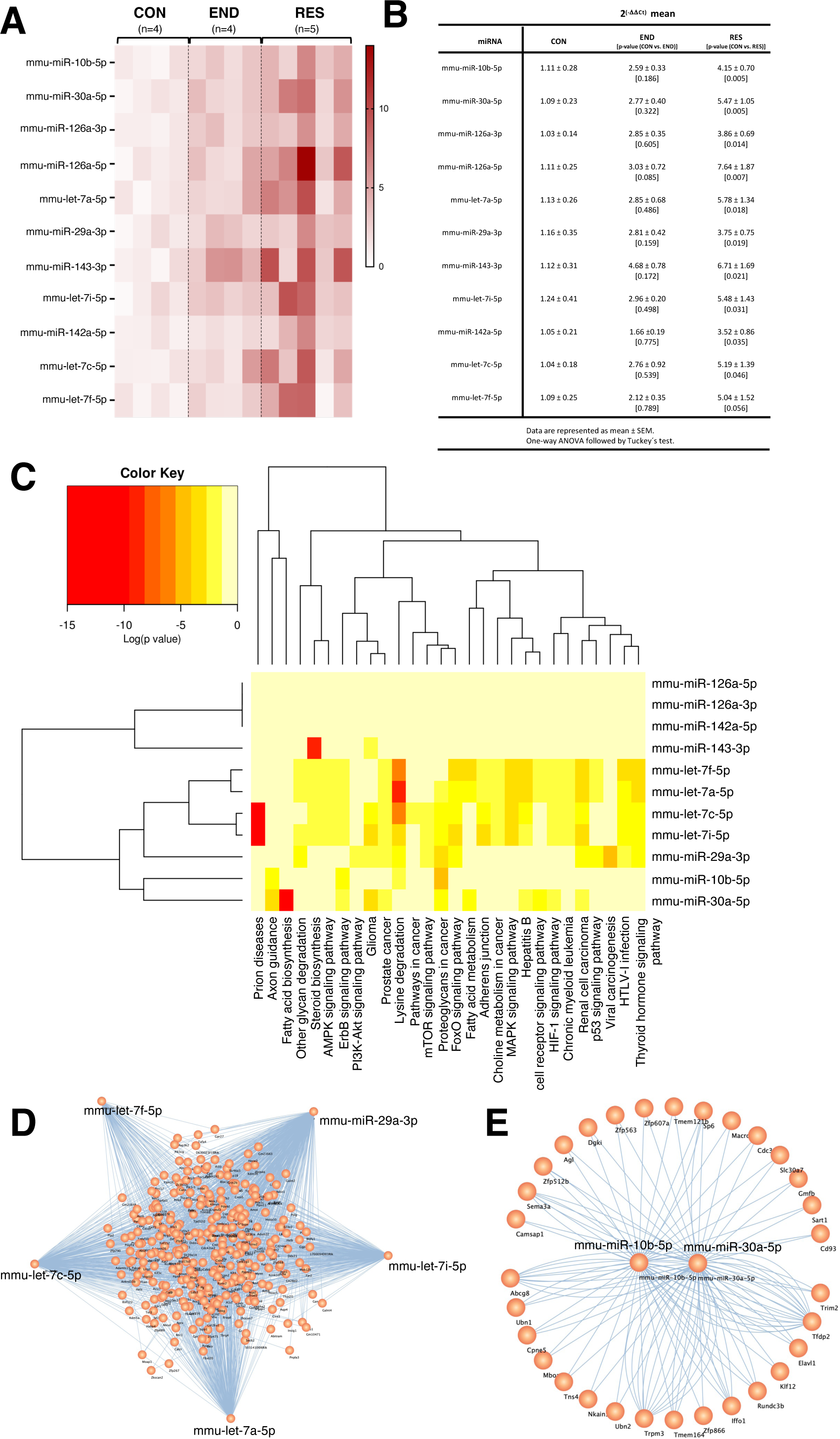
Training-associated miRNA profile in extracellular vesicles (EVs), main KEEG pathways and functional miRNA clusters. (A) Heat map of the differentially expressed miRNAs identified by NGS and validated by RT-qPCR in CON (n=4), END (n=4) and RES (n=5). Each row represents one miRNA, and each column represents an individual mouse within each group. The expression levels are shown in colors, from white (low) to dark red (high). (B) Representative table of the relative expression (mean ± SEM) and p-value (CON vs. END) and (CON vs. RES) of the 11 training-associated miRNAs. Statistical significance measured by one-way ANOVA followed by Tuckey’s test. (C) Heat map of the pathways union analysis based on the validated targets of the 11 training-associated miRNAs, grouped by functional clusters. The heat map represents as logarithm (p-value) the probability of interaction of each miRNA with a specific pathway in *Mus musculus.* Significance levels are shown in different colors, ranging from yellow (low) to red (high). (D, E) Cytoscape representation of miRWalk gene regulatory networks of the miRNAs belonging to clusters 1 (D) and 2 (E) in *Mus musculus*.

Interestingly, significant correlations between the expression levels of these miRNAs were observed. Thus, regardless of exercise modality, let-7a-5p was positively correlated with let-7c-5p (R=0.96, END; R=0.91, RES) (Figure S2A and S2B). Moreover, specific correlations dependent on exercise modality were also observed. In the case of END (Figure S2A), positive correlations were found between miR-126a-5p with let-7f-5p (R=0.95); miR-126a-3p with let-7i-5p (R=0.99); and miR-10b-5p with miR-142a-5p (R=0.96). Meanwhile, miR-143-3p correlated negatively with miR-30a-5p (R=-0.98). RES training showed positive correlations between let-7f-5p with let-7i-5p (R=0.92) and with miR-30a-5p (R=0.90); and between miR-10b-5p with miR-126a-3p (R=0.90) and miR-29a-3p (R=0.94) (Figure S2B). In the case of the CON, positive correlations were observed between circulating levels of expression of various miRNAs (Figure S2C), but none of the associations matched those observed in the training groups.

Then, we performed an in-silico analysis, using DIANA-miRPath v.3.0, to identify the main integrated metabolic pathways in which the 11 miRNAs are involved (Figure 2C). This analysis revealed several processes associated with the response to exercise, such as central fat metabolism, fatty acid and steroid biosynthesis, and the AMPK, PI3K-Akt and mTOR signaling pathways. In addition, this analysis allowed us to identify two functional clusters. Thus, cluster 1 consists of let-7a-5p, let-7c-5p, let-7i-5p, let-7f-5p and miR-29a-3p, with 286 common targets (Figure 2D), and cluster 2, which includes miR-10b-5p and miR-30a-5p, with 30 common targets (Figure 2E).

Taken together, these results show that training, regardless of exercise modality, exerts a common EV miRNA signature, with a greater response to resistance exercise. The existence of two functional clusters suggests a coordinated response of these EV miRNAs for the physiological modulation of energy metabolism in the adaptation to training.

### Modulation of the EV miRNA signature in skeletal muscle in response to training suggests a role for EV miRNA trafficking during exercise

To better understand whether skeletal muscle is involved in tissue-to-tissue communication in response to training, through miRNA release as EV cargo, we next evaluated the expression levels of the miRNA signature identified, in two different types of skeletal muscles, soleus and quadriceps.

In the case of soleus, most of the training-associated miRNAs showed lower expression levels in both END and RES compared to CON. This is the case for let-7a-5p, let-7i-5p and miR-29a-3p, from cluster 1 (Figure 3A), miR-10b-5p and miR-30a-5p belonging to cluster 2 (Figure 3B), and two non-clustered miRNAs, miR-126a-5p and miR-126a-3p (Figure 3C). It is noteworthy the lower expression of let-7c-5p and miR-142a-5p and higher of let-7f-5p only in response to endurance training (Figures 3A and 3C). To analyze the relationship between the miRNA circulating and tissue levels, a Pearson correlation was performed. In the case of let-7c-5p, a negative correlation was observed in both END (R=-0.96; p=0.039, Figure S3A) and RES (R=-0.82; p=0.085, Figure S3B). In a training-specific manner, let-7f-5p showed a negative correlation in END (R=-0.93; p=0.075; Figure S3A), whereas miR-143-3p displayed a negative correlation in RES (R=-0.82; p=0.090, Figure S3B).

**Figure 3.**
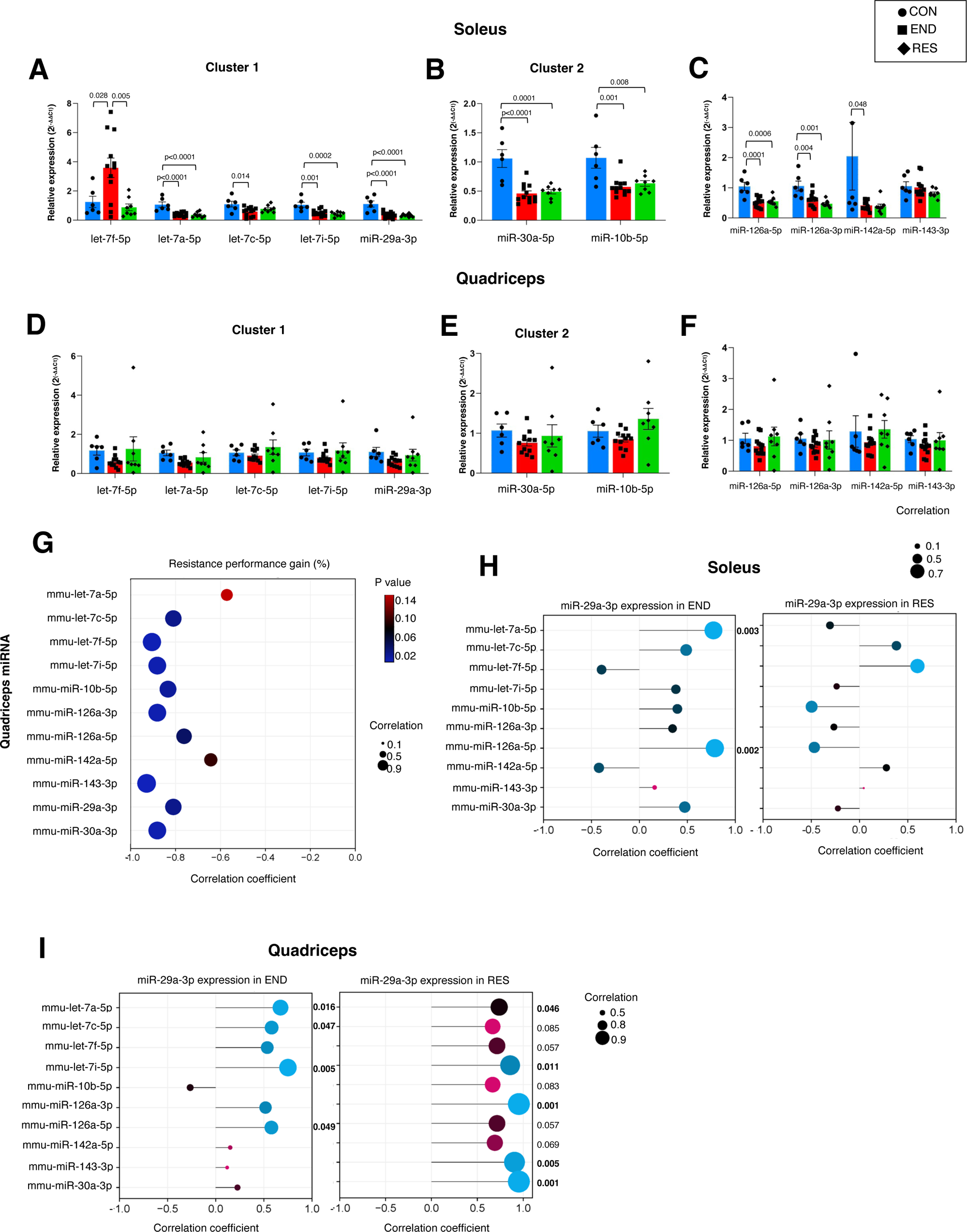
Modulation of the EV miRNA signature in skeletal muscle in response to endurance and resistance training. (A-C) Relative expression levels of miRNAs from clusters 1 (A), 2 (B) and unclustered miRNAs (C) in the soleus of CON, END and RES. (D-F) Relative expression levels of miRNAs from clusters 1 (D), 2 (E) and unclustered miRNAs (F) in the quadriceps of CON, END and RES. (G) Correlation of miRNA expression in quadriceps of RES and performance gain (%). (H) Correlation between miRNA expression in soleus of END and RES. (I) Correlation between miRNA expression in quadriceps of END and RES. Data are presented as mean ± SEM. Each dot represents one mouse (CON, n=6; END, n=12 and RES, n=8). Statistical significance measured by one-way ANOVA followed by Tuckey’s test.

In contrast, none of the miRNAs analyzed showed an altered expression in quadriceps (Figure 3D-F). However, Pearson correlation analysis referring to plasma versus quadriceps expression levels revealed a positive correlation for miR-29a-3p in END (R=0.97; p=0.035, Figure S3C) and RES (R=0.86; p=0.059, Figure S3D), while miR-10b-5p showed a negative correlation only in END (R=-0.96; p=0.048, Figure S3C). In addition to the correlation observed between plasma and quadriceps expression levels, the expression of these two miRNAs in this muscle tissue together with the expression of let-7c-5p (R=-0.81, p=0.022), let-7f-5p (R=-0.91, p=0.005), let-7i-5p (R=-0.88, p=0.005), miR-126a-3p (R=-0.88, p=0.007), miR-143-3p (R=-0.93, p=0.002) and miR-30a-5p (R=-0.88, p=0.007) were negatively correlated with resistance improvement (Figure 3G), again highlighting the coordinated response to training of these miRNAs. This coordinated response was also evident in the quadriceps of END and RES, emphasizing miR-29a-3p as a pivotal orchestrator of the miRNA response in this context. Indeed, its levels significantly and positively correlated with let-7a-5p and let-7i-5p in both trained groups and with miR-126-3p, miR-143-3p, and miR-30a-5p in RES group. Moreover, in the RES group, a tendency (p<0.1) was observed between miR-29a-3p and the remaining training-associated miRNAs (Figure 3H-I).

In this sense, Figure S3E shows a strong significant positive correlation in quadriceps of RES for all training-associated miRNAs, except for miR-142a-5p, which was not expressed in muscle cells. This quite evident positive association exerted by resistance exercise on the quadriceps was also observed for the rest of the training-associated miRNAs, although it was weaker in the case of miR-142a-5p (Figure S3E).

In summary, EV miRNAs were also detected in skeletal muscle of trained and untrained animals, suggesting their secretory role, which we will study further by means of in vitro approaches. Moreover, their response to training was different according to the training model and muscle type and seems to be coordinated, highlighting the role of resistance training as a coordinator of training-associated miRNAs in the quadriceps, with miR-29a-3p as the cornerstone. This, coupled with the fact that the relationship between circulating and intracellular miRNA levels in soleus and quadriceps was heterogeneous, suggests a muscle type-specific secretory profile.

### MiR-29a-3p is secreted in EVs by mouse myotubes in response to electrostimulation and has a modulatory effect on the expression of training-associated miRNAs

To determine the possibility that miRNAs present in circulating EVs were secreted by skeletal muscle cells, we used the mouse myoblast cell line C2C12, which expresses contractile proteins associated with type II muscle fibers, and has been described as one of the most suitable models for exercise response studies ^16^. After differentiation of myoblasts to myotubes, they were maintained for 3 h with or without electrical pulse stimulation (EPS) to induce contractions, as a simulation of exercise ^16^. Then, both myotubes and culture media were collected at three different time points, 0, 3, and 20 h after EPS, to isolate EVs (Figure S4A) for miRNA determination (Figure 4A).

**Figure 4.**
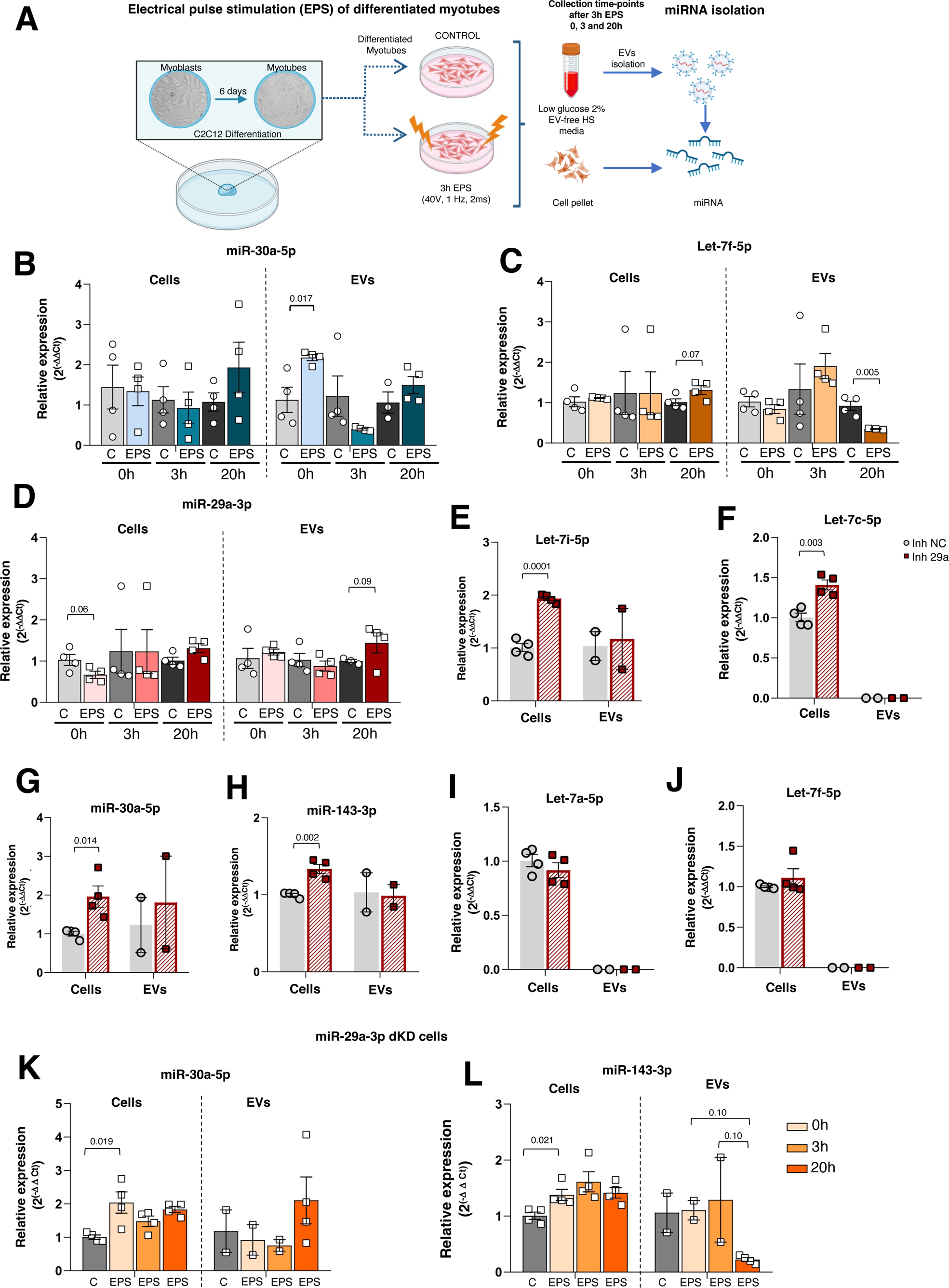
In vitro study of the impact of EPS in miRNA secretion and the modulatory effect of miR-29a-3p inhibition in training-associated miRNAs in mouse myotubes. (A) Schematic representation of electrostimulation protocol on C2C12 cells and isolation of miRNAs from EVs and cells. (B-D) Relative expression levels of miR-30a-5p (B), let-7f-5p (C), and miR-29a-3p (D) in cells and EVs at 0, 3, and 20 hours after electrostimulation in non-electrostimulated (C, n=4) and electrostimulated (EPS, n=4) cells. (E-H) Relative expression of miR-29 family members (E), cluster 1 miRNAs (F), cluster 2 miRNAs (G), and miR-143-3p (H) in C2C12 myotubes where miR-29a-3p expression was inhibited by 70% (miR-29a-3p dKD, n=4) and in control cells (NC, n=4). (I and J) Relative expression of miR-30a-5p and miR-143-3p in non-electrostimulated (C, n=4) and electrostimulated (EPS, n=4) miR-29a-3p dKD cells at 0, 3, and 20 hours after electrostimulation. Data are represented as mean ± SEM. Each dot represents the mean technical duplicate of each cell replicate. Statistical significance was determined using Student’s t-test for independent samples.

Most of the training-associated miRNAs identified in plasma EVs and skeletal muscle were also detected in the in vitro model (Figures 4B-D and S4B-E). On the contrary, miR-126a-3p, miR-126a-5p, miR-142a-5p, and miR-10b-5p were not detected, which may indicate that the main source of these four miRNAs is not of muscle cell origin. Therefore, the decreased levels observed in mouse muscle may be due to another cell types present in this tissue. Indeed, miR-126a-3p and miR-126a-5p are mainly expressed in endothelial cells ^17^, the main mononuclear cell type in muscle, but not in myotubes ^18^. In the case of miR-142a-5p and miR-10b-5p, their absence in the C2C12 cells had already been observed by other authors ^19^.

Of the miRNAs analyzed that showed expression changes in EPS C2C12 cells compared to unstimulated cells (Figure 4B-D), all of them displayed different dynamics. Thus, miR-30a-5p increased its expression in EVs recovered from the culture media immediately after the end of the EPS period (EVs, Time 0 h; Figure 4B). In contrast, the response of let-7f-5p was more delayed, as it decreased in EVs and tended to increase at the intracellular level 20 h after the end of EPS (Figure 4C). Finally, miR-29a-3p dynamics showed a tendency to be more spaced in time, as its expression was lower in EPS myotubes at time 0 and higher in EVs 20 h after EPS (Figure 4D). This miR-29a-3p response is similar to what we have observed in vivo, as its expression in plasma EVs was increased 24 h after the last training session (Figure 2A-B) and decreased in soleus (Figures 3B) in trained mice. Interestingly, this miRNA has been implicated in the glucose and fatty acid metabolism, two of the main fuels during exercise ^20,21^.

Next, we performed a downregulation study in C2C12 cells using a miR-29a-3p inhibitor that specifically binds to the target miRNA (miR29dKD cells; Figure S5A). First, we confirmed the inhibition of miR-29a-3p and the decrease in the expression of the other family members, miR-29b-3p and miR-29c-3p (Figure S4B), due to sequence similarities ^22^.

We first tested whether miR-29a-3p downregulation modulated the expression of the other training-associated miRNAs both intracellularly and in EVs. At intracellular level, its repression increased the expression of let-7i-5p and let-7c-5p, from functional group 1 (Figure 4E and 4F), miR-30a-5p, from group 2 (Figure 4G), and the unclustered miR-143-3p (Figure 4H). Interestingly, in relation to the encapsulation and export of these miRNAs in EVs, inhibition of miR-29a-3p did not appear to affect either let-7i-5p, miR-30a-5p or miR-143-3p. Interestingly, let-7c-5p expression was not detected in EVs, despite its high intracellular levels (Figure 4F).

Regarding the effect of electrostimulation in the absence of miR-29a-3p, EPS did not reverse the intracellular downregulation of the miR-29 family (Figures S5C and S5D). However, it appears to have a modulatory effect on miR-30a-5p and miR-143-3p by further increasing their expression at the intracellular level (Figure 4K and 4L). Regarding EV secretion, interestingly, the absence of miR-29a-3p appeared to prevent miR-30a-5p export in EVs after EPS (Figure 4K), in contrast to the secretion dynamics observed for this miRNA in non-inhibited cells (Figure 4B). Regarding miR-143-3p, while in the presence of miR-29a-3p it did not appear to respond to EPS, in the absence of this miRNA, it appears to be a tendency that its secretion in EVs was compromised at 20h post EPS. This inhibition of secretion of other miRNAs in the absence of *miR-29a/b1* can be seen in a more severe manner in the case of let-7f-5p, which was not detected in EVs (Figure 4J).

These results show that miR-29a-3p and miR-30a-5p are secreted in vitro by skeletal muscle in EVs in response to electrostimulation, although with different dynamics. The fact that in the absence of miR-29a-3p the intracellular expression of miR-30a-5p and miR-143-3p increased further after electrostimulation but decreased in EVs suggests a coordinated response to the contractile stimulus affecting secretion in EVs from members of different functional groups.

### Decreased physical capacity, particularly resistance, in *miR-29a/b1*-deficient mice might be related to dysregulated expression of training-associated miRNAs in quadriceps and EVs

From the in vitro results we have seen that miR-29a-3p is an myokine secreted by skeletal muscle. This is why we set out to analyze its functional role using a systemic *miR-29a/b1* cluster knock-out mouse (miR-29KO). This model is characterized by a life expectancy of ∼30 weeks, with a degenerative phenotype starting at 16 weeks of age ^20^. The lack of expression of miR-29a-3p was confirmed in EVs, quadriceps and soleus tissues (Figure 5A). Furthermore, the expression levels of miR-29b-3p were drastically reduced in plasma EVs and both skeletal muscles (Figure 5A). This indicates that miR-29b1 is the main form of miR-29b-3p that is expressed in these structures. However, miR-29c-3p only showed a decrease in soleus, with no changes in EVs and quadriceps (Figure 5A).

**Figure 5.**
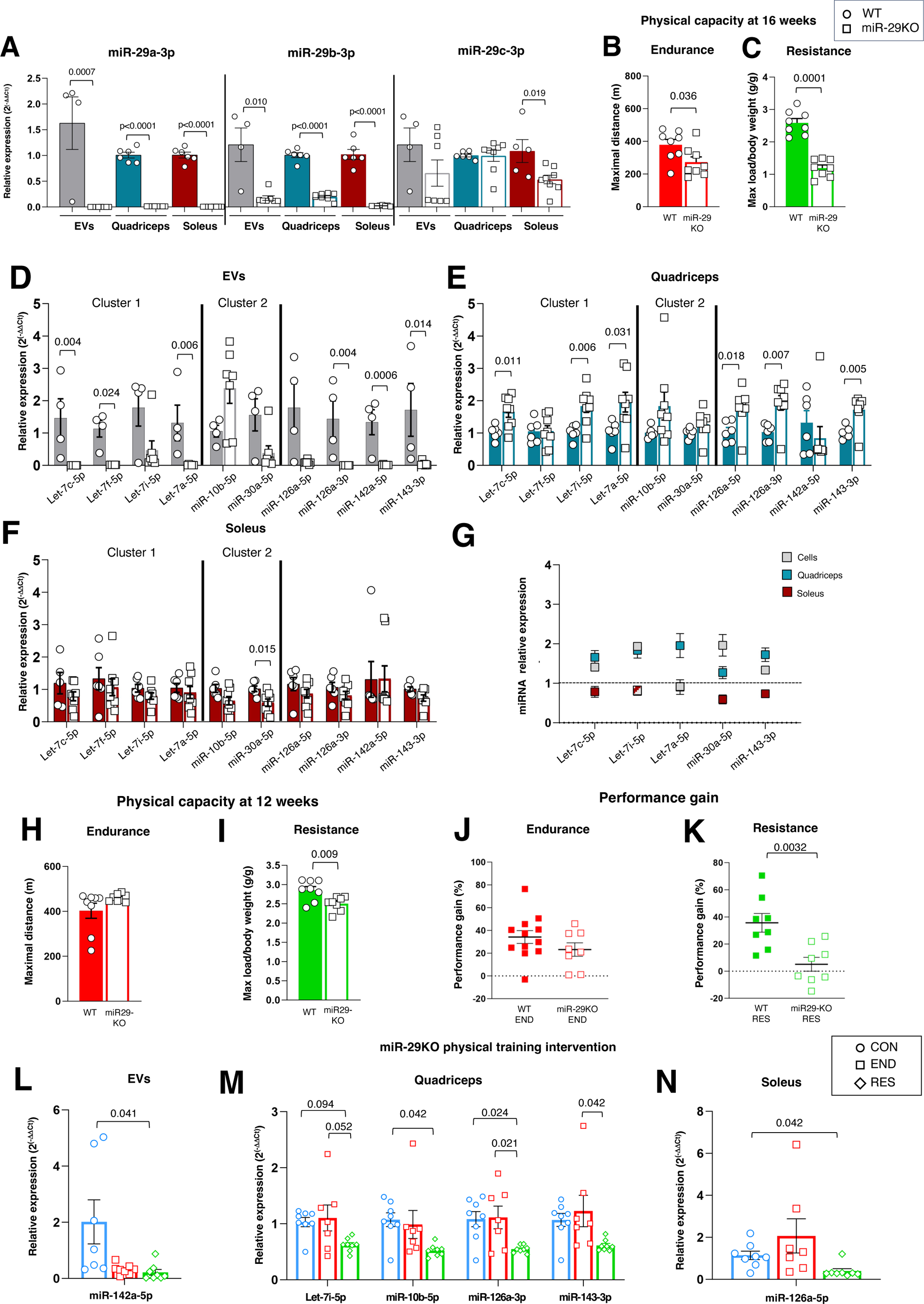
*MiR-29a/b1* knockout mice show low resistance capacity and an altered training-associated miRNA profile in skeletal muscle. (A) Relative expression levels of miR-29 family members in EVs, quadriceps, and soleus of 16-week WT and miR-29KO mice. WT (n=4 EVs, n=6 skeletal muscle tissue), KO (n=8 for all data). (B) Endurance performance of 16-week WT and miR-29KO mice (WT, n=8; miR-29KO, n=8). (C) Resistance performance of 16-week WT and miR-29KO mice (WT, n=8; miR-29KO, n=8). (D-F) Relative expression levels of the 11 training-associated miRNAs organized by clusters in EVs (D), quadriceps (E), and soleus (F) of 16-week WT and miR-29KO mice. (H) Relative miRNA expression in quadriceps and soleus of miR-29KO mice versus miR-29a-3p dDK myotubes differentiated in vitro (n=8 independent samples of tissue and n=4 independent samples of cell culture). (I) Endurance performance of 12-week WT and miR-29KO mice (WT, n=8; miR-29KO, n=8). (J) Resistance performance of 12-week WT and miR-29KO mice (WT, n=8; miR-29KO, n=8). (K) Endurance performance gain measured as percent variation of END and miR-29KOEND (END, n=12; miR-29KOEND, n=8). (L) Resistance performance gain measured as percent variation of RES and miR-29KORES (RES, n=8; miR-29KORES, n=8). (M-O) Relative expression levels of altered miRNAs after training, in EVs (M), quadriceps (N) and soleus (O) of miR-29KOCON (n=8), miR-29KOEND (n=7), and miR-29KORES (n=8). Data are presented as mean ± SEM. Each dot represents one mouse. Statistical analysis measured by Student’s t-test for all analysis except for Figures M, N and O, where statistical significance was measured by one-way ANOVA followed by Tuckey’s test.

Thus, we evaluated both endurance and resistance physical capacity in a group of 16-week-old miR-29KO mice and their WT counterparts. Although both capacities were affected, this effect was greater in resistance (53% mean loss) than in endurance (25%) (Figures 5B and 5C). This indicates that *miR-29a/b1* cluster plays a more critical role in resistance than in endurance performance.

One of the reasons that could explain the role of *miR-29a/b1* cluster in physical capacity is that, as observed in C2C12 miR29dKD myotubes, its absence could have a modulatory effect on other training-associated miRNAs at the systemic level. Accordingly, these miRNAs showed a dramatic reduction in their expression levels in EVs of miR-29KO mice regarding WT, except miR-10b-5p (Figure 5D). In the case of skeletal muscle, the observed response appeared to be dependent on the type of muscle. Thus, an increased expression of let-7c-5p, let-7i-5p, let-7a-5p, miR-126a-5p, miR-126a-3p and miR-143-3p was detected in quadriceps of miR-29KO mice (Figure 5E). Meanwhile, in soleus, only miR-30a-5p presented lower expression levels (Figure 5F). These results show that quadriceps is more sensitive to *miR-29a/b1* cluster absence. Therefore, the lack of expression of the *miR-29a/b1* cluster influences other training-associated miRNAs in miR-29KO mice in this tissue, and this may be one of the reasons for the decrease in endurance and resistance capacity observed in these animals. In fact, the higher the dysregulation in quadriceps of let-7a-5p and miR-143-3p, the worse resistance and endurance performance, respectively (R=-0.60, p=0.11; R=-0.59, p=0.13).

To compare these results with those obtained in the in vitro model, we plotted the relative expression levels of those miRNAs whose expression changed significantly in the skeletal muscles of miR-29KO mice and were also detected in miR29dKD cells (Figure 5G). Overall, the in vitro model showed more similar behavior to the quadriceps, as observed for let-7c-5p, let-7i-5p, miR-30a-5p and miR-143-3p, while let-7a-5p was more similar to the soleus (Figure 5G).

### *MiR-29a/b1* cluster plays a role in muscle mass maintenance by Murf1-independent mechanisms

One of the main phenotypic characteristics of miR-29KO mice is muscle atrophy ^23^, which could justify the decreased physical capacity in this model at early ages compared to WT mice. Indeed, by micro-computed tomography (micro-CT) we observed an 8.8% lower muscle area of the right hind paw per body weight in miR-29KOCON mice compared to CON (Figure S6A). It has previously been described the role of the *miR-29a/b1* cluster in muscle atrophy, as it regulates the expression of muscle ring finger protein-1 (MuRF1) ^23^. To determine whether this was extensible to our model, we assessed the expression levels and the protein synthesis of *Trim63*/Murf1 in quadriceps and soleus of 16-week-old sedentary miR-29KOCON and WT mice. Although we detected an increase in the expression of this gene in soleus but not in quadriceps (Figure 6A), there were no changes at the protein level in neither of the muscles (Figure 6B, 6C). This could indicate that the presence of the other members of the cluster, *miR-29b2/c*, is sufficient to maintain *Trim63*/Murf1 levels, as previously described ^23^

**Figure 6.**
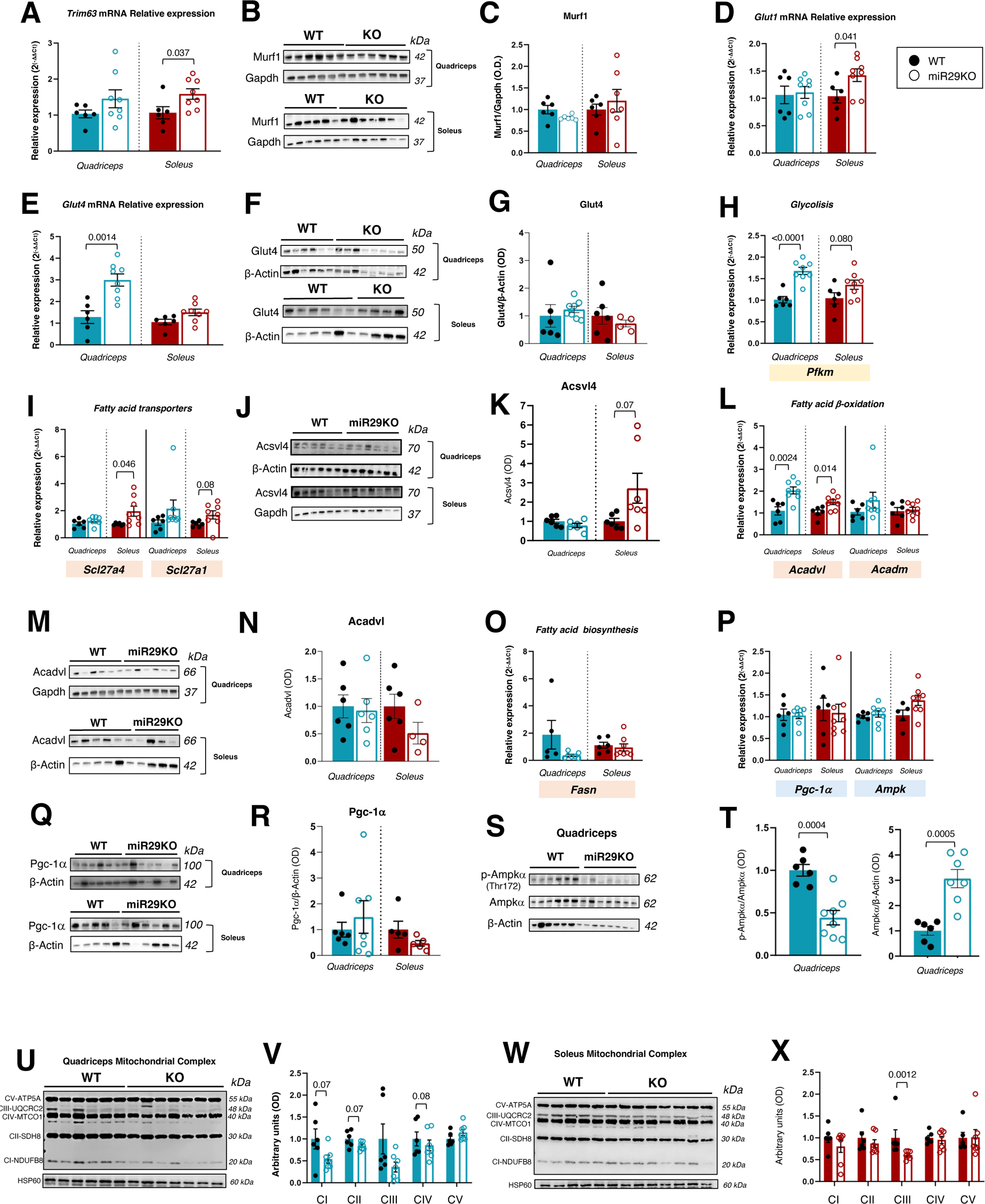
The absence of *miR-29a/b1* increases the availability of energy substrates in quadriceps and soleus but negatively affects their use. (A) *Trim63* relative expression in WT and miR-29KO mice. (B and C) Murf1 protein levels in WT (n=6) and miR-29KO (quadriceps, n=6; soleus, n=7). (D and E) *Glut1* and *Glut4* relative expression in WT and miR-29KO mice. (F and G) Glut4 protein levels in WT (n=6) and miR-29KO (quadriceps, n=8; soleus, n=7). (H) *Pfkm* relative expression in WT and miR-29KO mice. (I) *Slc27a4* and *Slc27a1* relative expression in WT and miR-29KO mice. (J-K) Acsvl4 protein levels in WT (n=6) and miR-29KO mice (quadriceps, n=8; soleus, n=7). (L) *Acadvl* and *Acadm* relative expression in WT and miR-29KO mice. (M-N) Acadvl protein levels in WT (n=6) and miR-29KO mice (quadriceps, n=6; soleus, n=4). (O) *Fasn* relative expression in WT and miR-29KO mice. (P) *Pgc-1*α and *Ampk* relative expression in WT and miR-29KO mice. (Q-R) Pgc-1α protein levels in WT (quadriceps, n=6; soleus, n=5) and miR-29KO mice (quadriceps, n=7; soleus, n=6). (S-T) p-Ampkα (Thr^172^) and total Ampkα protein levels in WT (n=6) and miR-29KO mice (n=8). (M-P) Mitochondrial respiratory complex OXPHOS protein levels in WT and miR-29 KO mice. Data are presented as mean ± SEM. Each dot represents one mouse. For mRNA analysis we used n=6 WT and n=8 miR-29KO mice and β-Actin and Gapdh as housekeeping genes. Statistical significance measured by Student’s t-test for independent samples.

The fact that this loss of muscle mass showed different impact on the resistance and endurance capacities of miR-29KO mice opens the door to consider other mechanisms than atrophy involved in this response, such us differences in energy metabolism, as suggested by functional analysis (Figure 2C).

### The absence of *miR-29a/b1* increases energy substrate bioavailability but not oxidative potential in muscle tissue leading to loss of physical performance

A shift in energy fuel utilization from fatty acid-based to glucose consumption-based metabolism was previously described in cardiac tissue of this murine miR-29KO model, with an increased expression of the ubiquitous glucose transporter *Glut1* ^20^. This exacerbation of glucose metabolism, related to the positive regulation of several key enzymes in this pathway, was also observed in the liver tissue of these miR-29KO mice ^20^. Therefore, we analyzed the expression of this transporter and detected that *Glut1* levels were higher in soleus (Figure 6D) and liver (Figure S7A) of miR-29KO mice compared to their WT counterparts, but not in quadriceps (Figure 6D). Interestingly, the inducible transporter *Glut4* showed higher expression levels in the quadriceps of miR-29KO, but not in soleus (Figure 6E), although this was not accompanied by changes at protein level (Figures 6F, G). We also determined the expression of the main regulatory enzyme of the glycolytic pathway, phosphofructo-1-kinase (*Pfkm*). *Pfkm* expression was higher in both muscle tissues of miR-29KO mice (Figure 6H). However, in the case of the soleus, although a trend was observed, the difference was not significant. Thus, it appears that glucose transport and utilization in skeletal muscle is regulated differently in the absence of *miR-29a/b1*, consistent with that previously described for cardiac muscle and liver ^20^.

In order to analyze the functional relevance of these differences, we conducted a maximal incremental endurance test in other cohort of 16-week-old miR-29KO (n=4) and their corresponding WT (n=4) mice, in which blood lactate and glucose levels were determined every 3 min during the test. Results showed no differences in basal lactate levels between genotypes. Furthermore, lactate levels in miR-29KO mice remained stable during the test, while a significant 65% increase (or 1.7±0.8 mmol/L) was observed in WT mice (Figure S7F). In line with what was previously described by Ferreira et al.^24^, this increase was not linear, and lactate accumulation started at 40 cm/s, reaching 4.3±0.6 mmol/L. No changes in glucose levels were observed during the test in miR-29KO mice, while there was a tendency in WT mice (12% increase or 22±16 mg/dL) (Figure S7G). This suggests that *miR-29a/b1* cluster deficiency could be affecting the transition from aerobic to anaerobic metabolism, due to a differential profile of substrate use.

The low blood lactate levels measured in miR-29KO mice may indicate that it is being used as a gluconeogenic substrate at the hepatic level. Indeed, higher liver expression of fructose-1,6-bisphosphatase 1 (*Fbp1*), an important enzyme of gluconeogenesis, was observed in this tissue in miR-29KO mice compared to WT (Figure S7A).

When analyzing genes involved in fatty acid metabolism, we observed that *Scl27a4* and *Scl27a1* fatty acid transporters were overexpressed in soleus and liver of miR-29KO mice (Figure 6I and Figure S7B), with a tendency to higher protein levels of Acsvl4 in soleus (Figure 6J and Figure 6K). Interestingly, no differences in mRNA or protein levels were observed in quadriceps (Figure 6I, Figure 6J and Figure 6K). Regarding β-oxidation, *Acadvl*, a very long-chain acyl-CoA dehydrogenase, and *Acadm*, a medium-chain acyl-CoA dehydrogenase, were studied. *Acadvl* gene presents higher levels of expression in skeletal muscle and liver in miR-29KO mice (Figure 6L and Figure S7C), although we have not found changes at the protein level in muscle tissue (Figure 6M and Figure 6N), while *Acadm* showed higher levels in the liver of miR-29KO mice (Figure S7C), but not in the quadriceps or soleus (Figure 6L). Interestingly, no changes in the expression of fatty acid synthase, *Fasn*, was detected (Figure 6O and Figure S7D). These results confirm the findings in liver of miR-29KO animals by Caravia et al.^20^. Contrary, the same authors described a decreased expression of *Acadvl* gene in the cardiac tissue of this same KO mice, which indicates a tissue-specific response of fatty acid metabolism in the absence of *miR-29a/b1*. Taken together, these findings suggest that in 16-week-old sedentary miR-29KO mice there is an exacerbation in the availability of energy substrates in skeletal muscle and liver compared to their WT counterparts. The observed response in the liver of these animals may be partially attributed to dysregulation of the other two miR-29 family members (Figure S7H) and other training-associated miRNAs (Figure S7I), showing an increased level of expression (Figure 5E).

We therefore went on to evaluate the status of the energy sensor par excellence, AMPK. In this case, *Ampk* showed higher levels in the liver of miR-29KO mice (Figure S7E) and a tendency to increase in soleus, but not in quadriceps (Figure 6P). However, interestingly, in quadriceps, despite higher total Ampk protein levels, the phosphorylated active form ^25^, decreased in miR-29KO mice, which may indicate a reduced activation of AMPK pathway (Figure 6S and Figure 6T).

Given these results, we decided to explore the expression of the *Pgc-1α* gene, one of the main targets of the miR-29 family ^20^, which is involved in the regulation of both carbohydrate and fatty acid metabolism. It was previously described that the expression of this gene was upregulated in the cardiac tissue and liver of more than 30-week-old miR-29KO mice^20^. However, at 16 weeks in the same model, our data showed no significant differences in either *Pgc-1α* gene expression or protein level in both skeletal muscles (Figure 6P, Figure 6Q and Figure 6R). Furthermore, in the liver we also observed no difference at the mRNA level (Figure S7E). This may indicate that this overexpression observed miR-29KO develops with aging.

Since this could be due to alterations in mitochondrial function, we analyzed protein levels of the 5 mitochondrial complexes in muscle tissues. The results showed that in the quadriceps there is a tendency for complexes I, II and IV to decrease (Figures 6M, N), while in the soleus complex III presents significant lower values in miR-29KO mice (Figures 6O, P). This suggests an underlying mitochondrial dysfunction particularly evident in quadriceps which could affect the effective use of energy substrates despite the increase observed in their availability, impairing a correct adaptation to acute exercise and training.

### *MiR-29a/b1* cluster has a profound effect on the adaptive response to resistance training in skeletal muscle

Since the absence of the *miR-29a/b1* cluster affects both the expression levels of training-associated miRNAs and energy metabolism genes at muscle level, influencing physical capacity, we wanted to study its impact on training adaptation.

For this purpose, we used 12-week-old miR-29KO mice and their corresponding WT controls. A 4-week training intervention, analogous to that previously described, was carried out at the same relative intensity.

The initial physical capacity test showed that miR-29KO mice had similar levels of endurance than CON (Figure 5H), while their resistance capacity was significantly lower (Figure 5I). This suggests that the loss of resistance observed at 16 weeks (Figure 5C) is already present at 12 weeks in this model.

After the training period, endurance-trained miR-29KO mice (miR-29KOEND) improved endurance performance similar to END (Figure 5J). However, resistance-trained mice (miR-29KORES) were only able to maintain their already impaired resistance capacity (Figure 5K). Thus, *miR29a/b1* cluster is required for the adaptive response to resistance but not to endurance training.

Regarding miRNA response, miR-142a-5p showed lower levels in EVs of miR-29KORES compared to miR-29KO sedentary mice (miR-29KOCON) (Figure 5L and Figure S8A). This miRNA was already downregulated in miR-29KOCON compared to CON (Figure 5D). No further changes were observed in any of the other miRNAs analyzed in plasma EVs. Therefore, the exercise intervention was ineffective in reversing the dysregulation observed in EV miRNAs of miR-29KOCON.

At the muscle level, a different response was observed between quadriceps and soleus, as well as between training interventions. Thus, resistance training induced a lower expression of let-7i-5p, miR-10b-5p, miR-126a-3p, and miR-143-3p (Figure 5M; Figure S8B) in quadriceps, while in soleus a lower expression of miR-126a-5p was observed (Figure 5N; Figure S8C), with no effect of endurance intervention. Therefore, resistance training intervention partially reversed the miRNA dysregulation observed in quadriceps of miR-29KOCON (Figure 5E), but not in soleus (Figure 5F). As for the liver, exercise had no effect on the modulation of miRNAs (Figure S9B).

At the muscle level, the improvement observed in miR-29KOEND mice could be caused by their 9.7% increasement in muscle mass compared to miR-29KOCON mice, reaching very similar levels than CON mice. However, for miR-29KORES an increase of 2.4% was observed, which is still 6.7% below CON animals (Figure S6B). Again, this process seems relevant to their functional capacity and adaptation to training and was independent of *Trim63* expression (Figure S6C and S6D).

As for the expression of genes related to energy metabolism (Figure S10), quadriceps was the main responsive tissue to training in miR-29KO animals in line with the changes observed in the expression levels of training-associated miRNAs (Figure 5M, Figure 5N, Figure S8B). Furthermore, in both cases, the greatest response is observed with resistance training. Thus, miR-29KORES mice showed lower expression levels of *Glut1* and *Glut4* in quadriceps compared to miR-29KOCON mice (Figure 7A). However, Glut4 protein levels were higher in miR-29KORES than in miR-29KOCON (Figure 7B, 7C). One possible explanation is that the inducible Glut4 transporter is recycled between stimuli, remaining in a reservoir of cytoplasmic vesicles ^26^. Thus, in miR-29KOCON mice, this process could be impaired in the quadriceps. However, resistance training-mediated stimulation could counteract possible deficiencies in either internalization or maintenance and re-exposure of Glut4, thereby reducing the need for increased *Glut4* expression. Similarly, the energy sensor *Ampk* exhibited lower levels of expression in miR-29KORES mice regarding miR-29KOCON (Figure 7D), whereas no changes in total protein levels were observed. However, the phosphorylated active form was higher than miR-29KOCON and miR-29KOEND (Figures 7E, 7F). Therefore, it might be possible that resistance training favors the Ampk pathway in the quadriceps of these miR-29KORES mice, thus enhancing glucose uptake, via Glut4, and its subsequent utilization ^26^. Analysis of the fatty acid transporter Acsvl4, at both the expression and protein levels (Figure 7G-I), suggests that improved glucose utilization might reduce the need for a high fatty acid uptake. This contrasts with what was observed in WT animals, where resistance training mainly increased the expression of genes related to fatty acid metabolism in soleus and liver (Figure S11B and Figure S11C), similar to what was observed for miRNAs (Figure S9A). However, in the quadriceps of these WT mice we did not observe notable changes, nor in response to resistance or endurance training (Figure S11A). That is, resistance training does not restore the miR-29KO phenotype to a metabolic profile more similar to WT animals. Regarding the effect of training in the protein levels of the 5 mitochondrial complexes in the quadriceps of miR-29KO trained animals, exercise does not seem to affect their protein levels (Figure S12).

**Figure 7.**
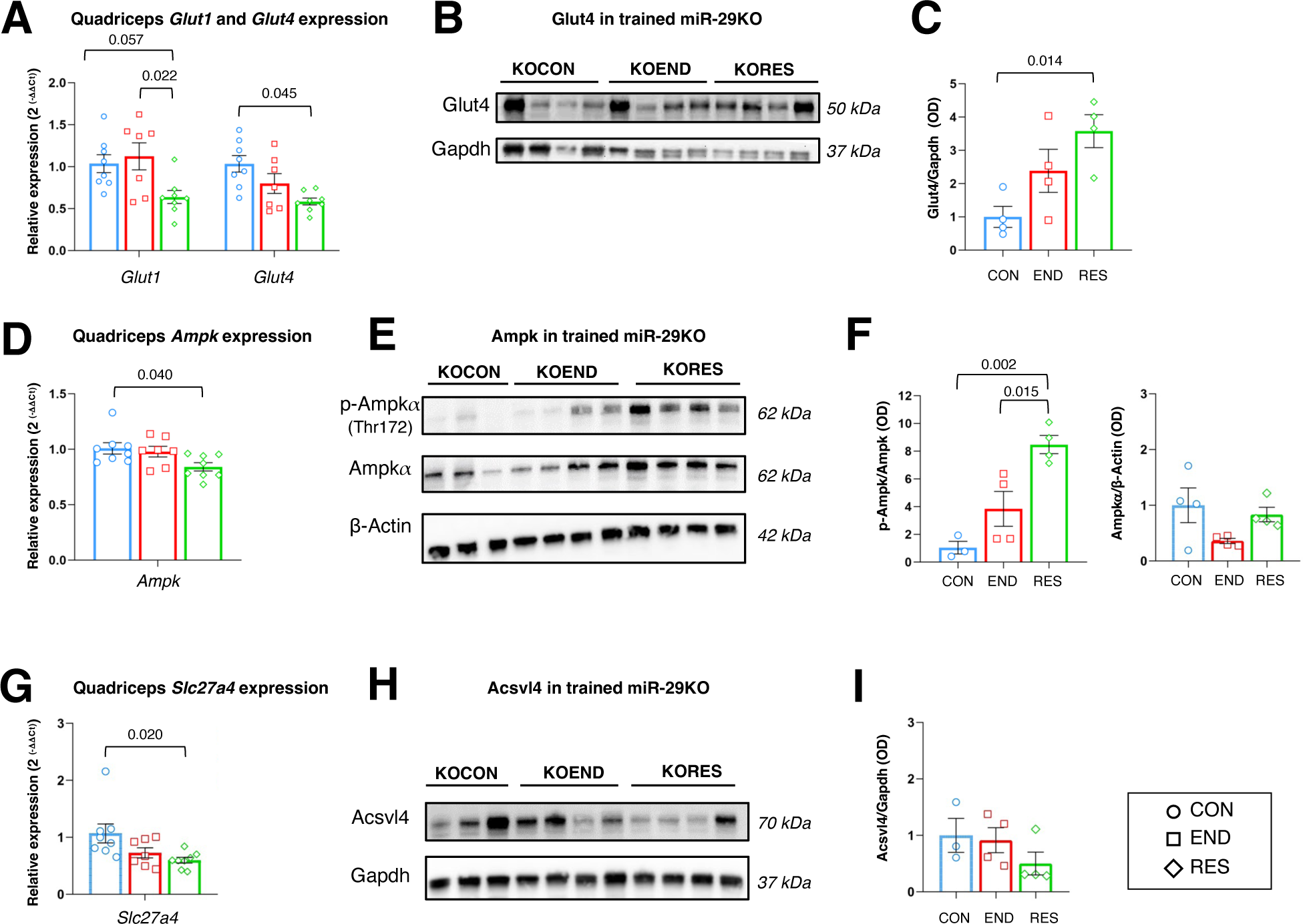
Resistance training exerts an effect on increasing glucose influx and energy production in the quadriceps of miR-29KO mice. (A) *Glut1* and *Glut4* relative expression in miR-29KOCON, miR-29KOEND and miR-29KORES. (B and C) Glut4 protein levels in the same groups (n=4/group). (D) *Ampk* relative expression in the same groups. (E and F) p-Ampkα (Thr^172^) and total Ampkα protein levels in the same groups (n=3 CON and n=4 END and RES). (G) *Slc27a4* relative expression in the same groups. (H and I) Acsvl4 protein levels in the same groups (n=3 CON and n=4 END and RES). Data are presented as mean ± SEM. Each dot represents one mouse. For mRNA analysis we used n=8 miR-29KOCON, n=7 miR-29KOEND and n=8 miR-29KORES. Statistical significance was tested by one-way ANOVA followed by Tuckey’s test.

Summarizing, *miR-29a/b1* cluster is essential for the adaptation to resistance training. However, resistance training halts the degeneration of resistance capacity that occurs in the absence of this cluster, suggesting that it is independent of *miR-29a/b1*. Muscle type-specific repression of other training miRNAs in response to resistance exercise may be mediating this process.

In summary, *miR-29a/b1* cluster is essential for adaptation to resistance training but independent of adaptation to endurance training. Specific repression of other training-associated miRNAs in the quadriceps in response to exercise coupled with prolonged activation of the Ampk/Glut4 axis favoring glucose uptake may be mediating this process.

## DISCUSSION

One of the mechanisms by which exercise mediates its beneficial effects consists of a systemic crosstalk mediated EVs containing different molecules, including miRNAs ^27 28 29^. However, it remains unresolved the secretory role of skeletal muscle in the response to training and which miRNAs encapsulated in EVs could be relevant in this context, acting as exerkines ^30^. Traditionally, the characterization of miRNAs contained in EVs in response to acute exercise and training has focused on skeletal muscle-associated miRNAs (myomiRs), such as miR-1, miR-206 and miR-133b (reviewed in ^7^). Castaño et al. ^15^ observed that most plasma EV miRNAs obtained from HIIT-trained mice were myomiRs, proposing skeletal muscle as the main secretory tissue of miRNAs encapsulated in EVs. However, recent research suggests that the presence of myomiRs is higher in interstitial EVs than in plasma EVs, making them more relevant in the context of skeletal muscle regeneration ^31^

Thanks to massive sequencing, we were able to determine that, predominantly resistance training, increases the expression of a group of 11 miRNAs in plasma EVs of mice in resting state, with no changes in myomiR expression. However, this does not rule out muscle as a secretory tissue for other miRNAs in response to training. In fact, our in vitro approaches support miR-29a-3p and miR-30a-5p as new myokines closely resembling the response observed in EVs extracted from the plasma of trained animals. Other authors have also reported the presence of these two miRNAs in EVs in response to an EPS protocol in C2C12 cells and in a more complex three-dimensional model called “myobundles”, respectively ^32,33^. The involvement of both miRNAs in muscle differentiation and metabolic function is well documented ^34–36^. Furthermore, we have observed that the inhibition of miR-29a-3p in cell cultures of myotubes increases the intracellular levels of miR-30a-5p and miR-143-3p, even more so after electrostimulation, yet seems to affect their export in EVs, which implies a synchronized response to the contractile stimulus among various miRNA from different functional clusters, involved in myogenesis and muscle cell differentiation ^34,36^. One possible cause of this coordination could be attributed to the role of miR-29a as an epimiRNA, referring to its function as a connector molecule for various epigenetic mechanisms^37^. In this sense, Rowlands et al. ^38^ supported the idea that, in response to endurance training, miR-29a downregulation in skeletal muscle in patients with type II diabetes was associated with the epigenetic regulation of the methylation process.

Interestingly, the in vitro model faithfully reflects this coordinated intracellular response, although only in the quadriceps. One possible explanation is that, as described by Abdelmoez et al.^16^ this C2C12 cell model showed a higher expression of MYH1 and MYH4 proteins associated with type II fibers, which are more predominant in the quadriceps ^39^. However, electrostimulation does not replicate the effect of resistance and endurance training on the intracellular expression of training-associated miRNAs in the muscles tested in miR-29KO mice, and in the case of miR-143-3p the response was the opposite. This difference could be due to our EPS protocol was more reflective of the acute response to endurance exercise, based on the frequency and time conditions used ^40^. Valero-Bretón et al.^41^ reported that replicating the acute and chronic stimulus effect of resistance exercise would have required increasing the stimulation frequency or the number of days, respectively, which was methodologically unfeasible in our model, due to the inhibition protocol used.

These results raise the question of whether the increase in miR-29a-3p observed in EVs our in vivo model could be due to a prolonged residual effect of the last exercise session, described by Darragh et al.^8^ as “last bout effect”, rather than the effect of chronic adaptation to training. However, despite limited information in rodent studies, Castaño et al.^15^ and Hou et al.^43^ described changes in miRNA profile in EVs taken at rest after a period of training. In addition, Whitham et al. ^4^ reported that in humans 4 hours after acute exercise EVs concentration returns to basal levels and previous data from our laboratory showed that several of the miRNAs observed in our study, including miR-29a-3p, increased at the circulating level after a marathon in highly trained amateur runners and returned to basal levels 24 hours later ^44^. Therefore, it seems that miR-29a-3p emerges as an myokine with coordinated expression and secretion with other miRNAs. Even more, our results show that it is essential for the maintenance of physical capacity and for the adaptation to training, particularly resistance. Indeed, Davidsen et al.^45^ reported that low responders’ males to a 12-week strength training programme had lower expression levels of miR-29a-3p and less muscle hypertrophy. This may be mediated by its regulatory role in the expression of miR-30a-5p and miR-143-3p in the quadriceps, having an impact on muscle development and differentiation and on energy metabolism in this tissue.

The exacerbated response of genes related to energy substrate availability and use in miR-29KO mice ^21,46,47^ supports the observation that these animals adapt to endurance training similarly to END, despite their final lower muscle mass. Furthermore, the fact that miR-29KOEND animals increase their muscle mass highlights the relevance of this parameter in adaptation to endurance training. Both mechanisms, relevant for endurance training adaptation, are thus independent of *miR-29a/b1*. Contrary, miR-29KO mice showed compromised resistance capacity and lower muscle mass. Specific training did not improve these characteristics, although prevented strength loss. This same response was previously described by us in a murine model with systemic partial autophagy deficiency ^14^, emphasizing resistance training as an optimal intervention for states of muscular frailty through a variety of underlying molecular mechanisms. Further analysis of this adaptive response to resistance training underlines the importance of miRNA coordinated expression for resistance training adaptation and resistance capacity maintenance. Thus, the repression of let-7i-5p, miR-126a-3p and miR-143-3p observed in miR-29KORES quadriceps contrasts with the overexpression of these miRNAs in miR-29KOCON. Regarding miR-143-3p, its downregulation after strength exercise may play a role in the upregulation of fast type IIa and IIb fibers in the quadriceps ^48^ by increasing the efficiency of this muscle in the exercise response. Moreover, this repression could be related to increased glucose uptake, mediated by the Glut4 receptor, followed by activation of Ampk for energy production, and contrasted with decreased utilization of lipid metabolism. This prolonged activation of the AMPK/GLUT4 axis could decrease 4E-BP1 phosphorylation^49^, a downstream component of mTOR signalling pathway, leading to decreased muscle protein synthesis and muscle mass gain in these miR-29KORES mice, affecting their adaptation to training. This might suggest that changes in miRNA expression in quadriceps allow this adaptation to resistance training and the maintenance of this capacity, even in the absence of *miR-29a/b1*.

Taken together, this study first to define miR-29a-3p as a new myokine with a determinant role in the response and adaptation to resistance training, mediating a miRNA coordinated response in muscle development and energy metabolism. Additionally, these results suggest that resistance training could be a useful and beneficial intervention for those diseases in which miR-29 family members are downregulated, such as pancreatic and renal fibrosis, osteoporosis ^50^, and even with a major impact on neurodegeneration ^51^, as described in Alzheimer’s disease ^52^.

## Acknowledgments

P.P-H. was supported by a contract associated with grant AYUD/2021/5134 from FICYT (Plan de Ciencia, Tecnología e Innovación 2018-2022 del Principado de Asturias), cofunded by the Fondo Europeo de Desarrollo Regional (FEDER), Spain. J.C.-S. was supported by a contract associated with grant AC20/00017 from Instituto de Salud Carlos III, Spain, cofounded by EuroNanoMed III (grant 20-0084). A.C.-V. was supported by a contract associated with grant AYUD/2021/57540 from FICYT/FEDER, Spain.

This work was supported by the Fundación Tatiana Pérez de Guzmán el Bueno (Convocatoria de ayudas a proyectos de investigación en neurociencia, edición 2020) to C.T.-Z. and E.I.-G and by the Spanish Ministerio de Economía y Competitividad (DEP2012-39262 and DEP2015-69980 P) to EI-G and BF-G.

## Sample Author Contributions

Conceptualization, B.F-G, C.T-Z and E.I-G; Methodology, C.T-Z and E.I-G; Investigation, P.P-H, M.F-S, D.P; J.C-S, A.C-V, S.D-R, and P.G; Software, M.F-S; Writing – Original Draft, P.P.H; Writing – Review & Editing, M.F-S, C.T-Z and E.I-G; Funding Acquisition, B.F-G, C.T-Z and E.I-G; Resources, D.R-V, X.C, J.Z, A.K, C.L-O, P.G-R, C.T-Z and E.I-G; Supervision, M.F-S, C.T-Z. and E.I-G.

## Conflicts of Interest

The authors declare no conflict of interest.

## STAR Methods

### Key resources table

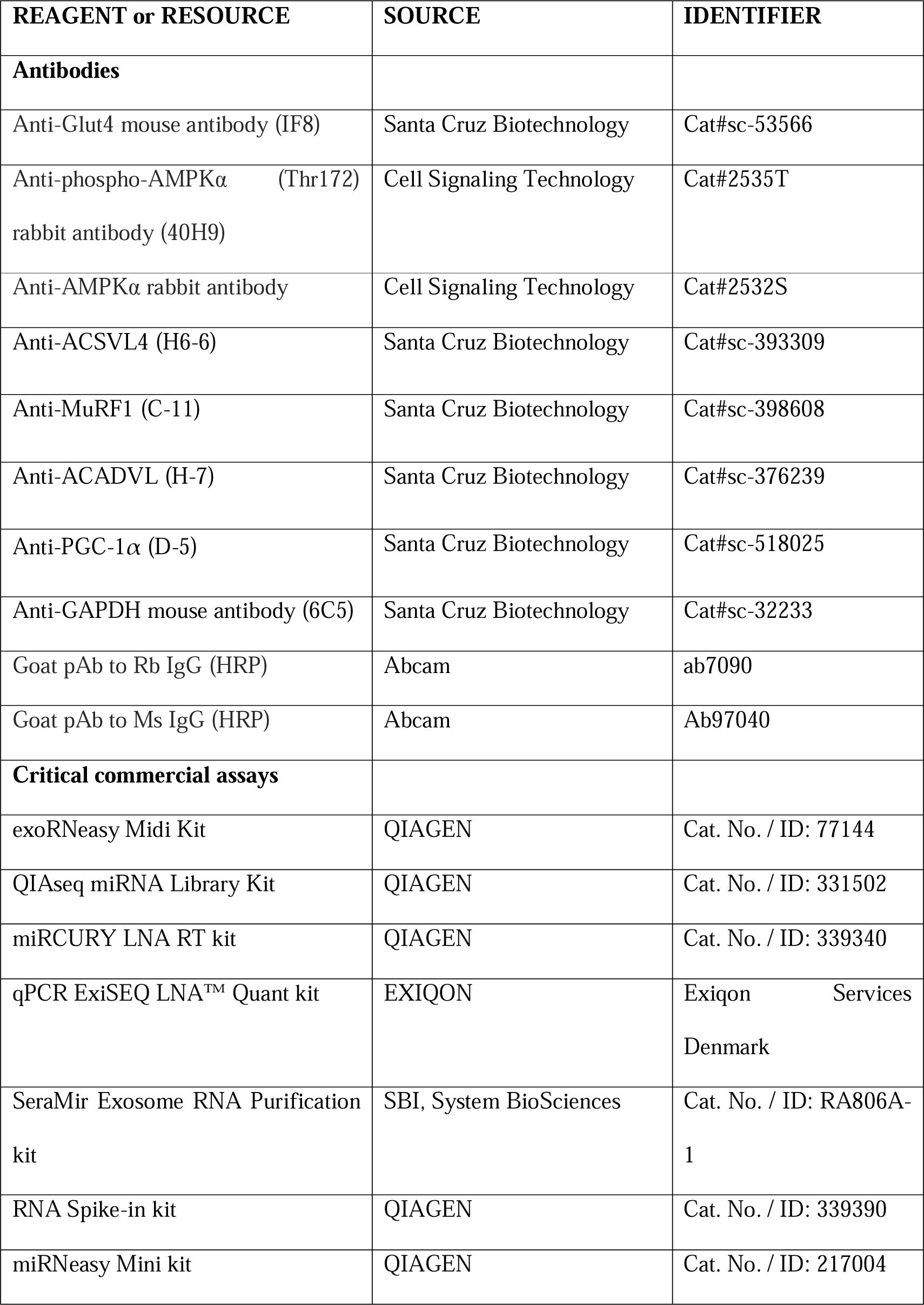
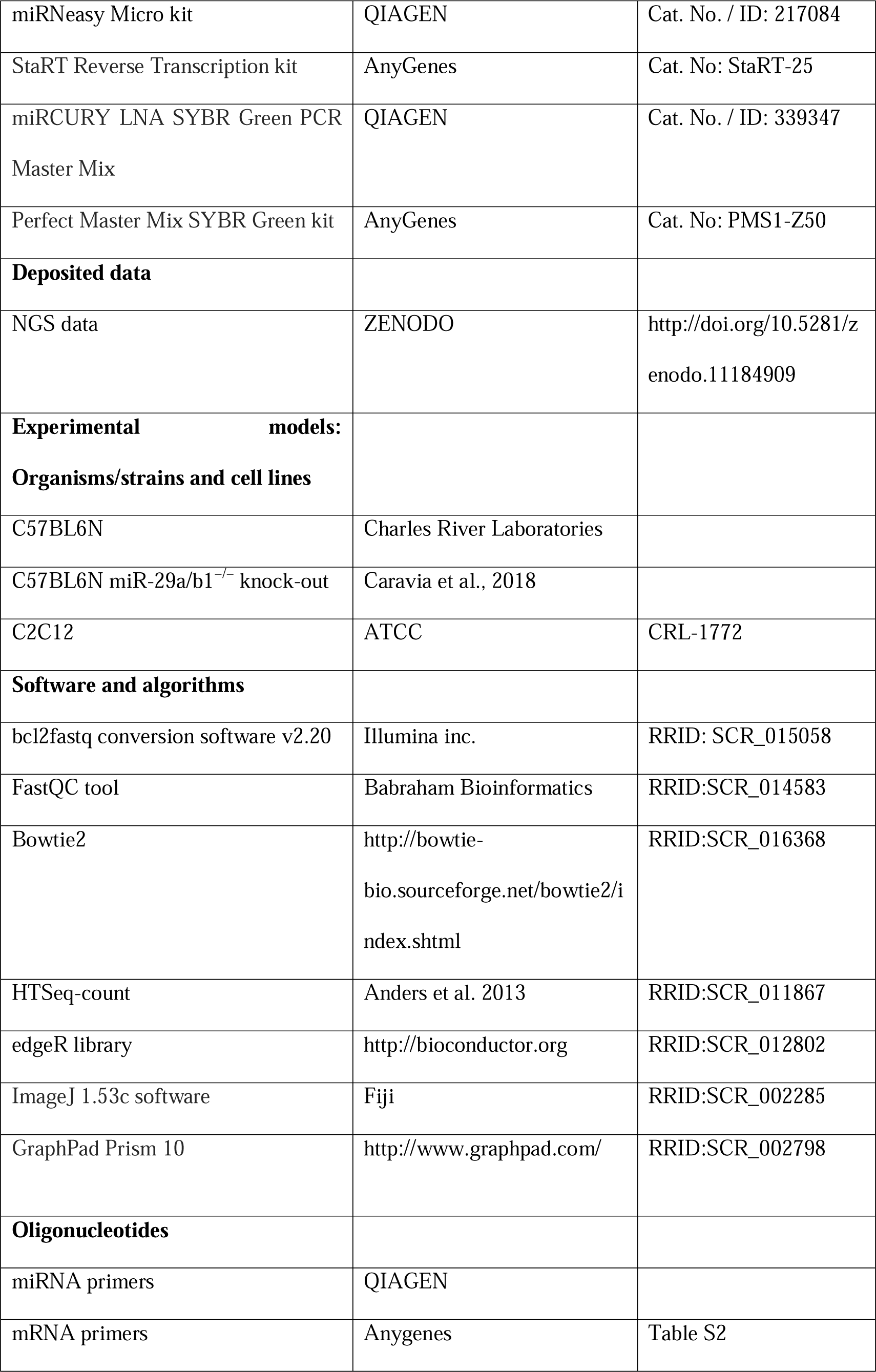
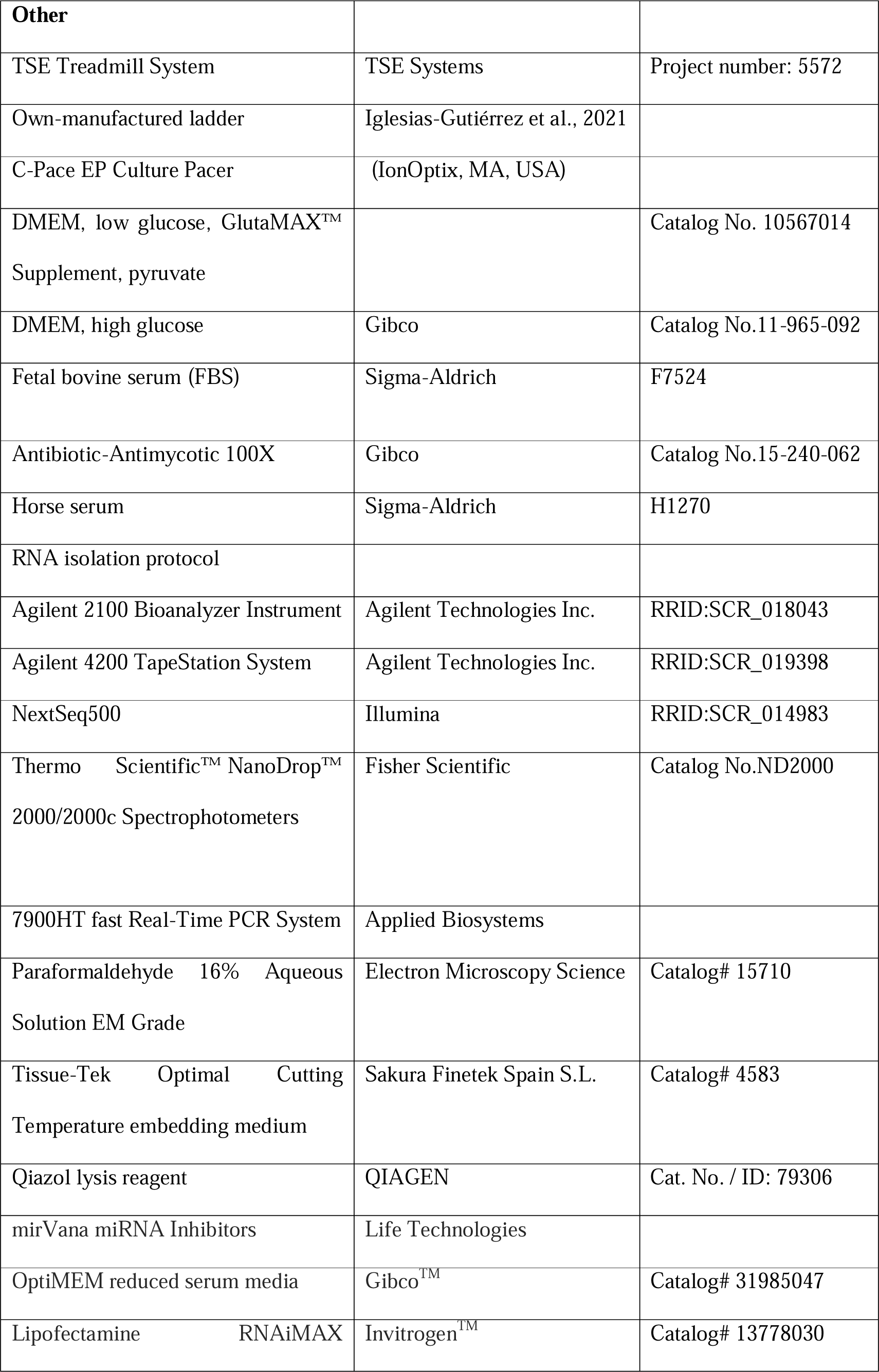
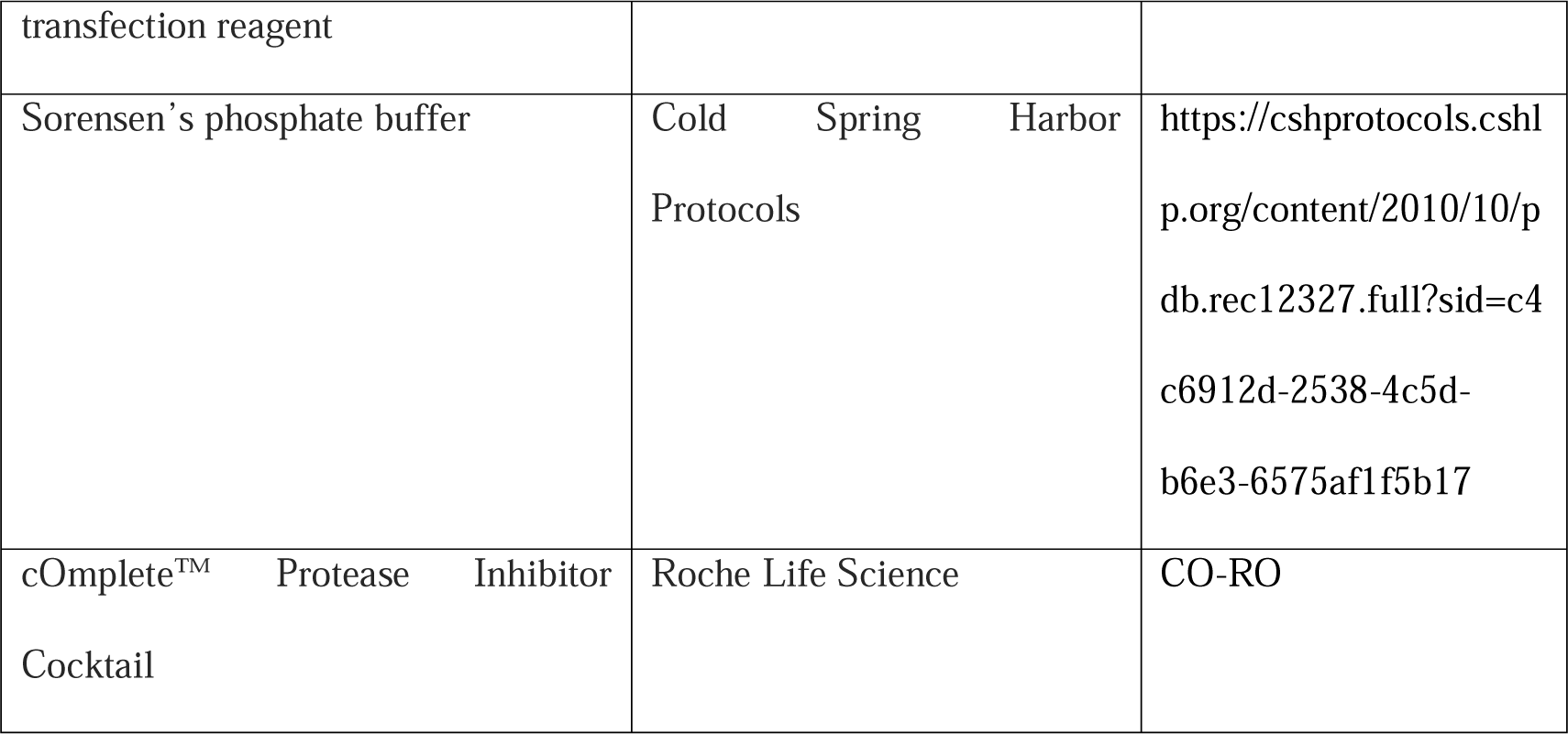

### Resource availability

#### Lead contact

Further information and requests for Data and resources should be directed to and will be fulfilled by the lead contact, Dr. Eduardo Iglesias-Gutiérrez.

#### Materials availability

This study did not generate new unique reagents.

### Experimental model and subject details

#### Animals

26 male mice of the same C57BL6 Wild-type (WT) genetic background were obtained from Charles River Laboratories. Male and female miR-29a/b1 heterozygous (miR-29a/b1^−/+^) mice of parental C57BL6 genetic background were provided by Dr. Carlos Otín’s research group ^20^. These mice were backcrossed with each other to obtain miR-29a/b1 knockout mice (miR-29a/b1^−/−^, miR-29KO). At one month of age, the offspring was genotyped and were separated according to genotype and sex.

A total of 26, 10-week-old, C57BL6N WT male mice and 23, 10-week-old, C57BL6N miR-29KO male mice were randomly divided into three different groups: sedentary control (CON, n=6; miR-29KOCON, n=8), endurance training (END, n=12; miR-29KOEND, n=7) and resistance training (RES, n=8; miR-29KORES, n=8).

Mice were fed using a pellet rodent diet (Teklad Irradiated Global 18% Protein Rodent Diet, Envigo, Spain) and water *ad libitum*. The food intake and the animals’ weight were measured weekly. Mice were maintained on a 12 h light/dark cycle (light onset at 8:00 AM) and under controlled temperature (22 ± 2 °C) at the Animal Facilities of the University of Oviedo, Spain (authorized facility No. ES330440003591). All procedures were conducted during the early light portion of the cycle and performed in accordance with the institutional guidelines approved by The Research Ethics Committee of the University of Oviedo, Spain (PROAE 10/2016 and PROAE 35/2020).

#### Cell culture of mouse C2C12 cells

The mouse skeletal muscle myoblast cell line C2C12 were obtained from the American Type Culture Collection (ATCC CRL-1772). The C2C12 cell line was cultured in DMEM (Dulbecco’s Modified Eagle Medium, 11965092, Gibco). This media contains high levels of glucose (4.5 g/L) and L-glutamine and was supplemented with 10% fetal bovine serum (FBS, F7524, Sigma-Aldrich) and 1% antibiotic-antimycotic cocktail (10000 U/mL penicillin, 10000 µg/mL streptomycin and 0.25 µg/mL amphotericin B (11580486, Gibco). For differentiation to myotubes, C2C12 myoblasts were seeded in a 6-well plate at a density of 2 × 10^6^ cells/well. After two days, when cells reached 90-100% confluence, differentiation was induced by switching to DMEM supplemented with 2% horse serum (HS, H1270-500ML, Sigma-Aldrich) and 1% antibiotic-antimycotic cocktail. Differentiation was monitored microscopically, and cells were used for experiments after 6-7 days. The absence of mycoplasma contamination was routinely confirmed by PCR. Cells were maintained at 37°C under 5% CO_2_.

### Method details

#### Animal physical training methods

A 4-week training intervention for WT mice and adapted with respect to the physical qualities of miR-29KO mice was performed. Two different types of training were performed, endurance and resistance training. Endurance exercise was performed on a treadmill without any aversive stimuli. A commercial four-rail treadmill (TSE Systems, Germany) with adjustable speed and incline was used. Resistance training was carried out in an own-manufactured static ladder ^53^. The ladder was built with 25 steel wire steps of 1.5 mm of diameter separated by 15 mm. A resting area of 20×20 cm was placed on top of the ladder The slope of the ladder was modifiable by the researcher, ranging between 90 and 80° with the horizontal plane. The same researcher handled and trained the mice during the different stages of training: acclimation period, specific maximal performance tests (pre- and post-training), and training protocols.

##### Acclimation period

Mice were acclimatized to the training devices for 2 weeks, 5 sessions per week, and 15 min per training session, as previously described in ^13,14^.

##### Maximal performance tests

Twenty-four hours after the end of the acclimation period, both WT and miR-29KO mice were randomly distributed in the above-mentioned training groups. Then, in the first week of training, maximal endurance and resistance tests were carried out. While training groups mice performed the corresponding maximal tests, sedentary mice performed both endurance and resistance tests 24 hours apart. Their maximal endurance capacity was determined by an incremental test in the treadmill and their maximal resistance capacity was tested in the vertical ladder, following the same protocols by our research group described before ^13,14^. Both tests were repeated at the end of the training period, following the same protocols.

Additionally, a maximal endurance test was carried out in a small cohort of 16-week-old WT (n=4) and miR-29KO (n=4) mice. Blood samples were taken from the tail vein at each step of the maximal exercise test (every 3 minutes) until exhaustion. At this point the animal was removed from the test and blood glucose and lactate levels were measured. The blood glucose concentration (mg/dL) was analyzed using Accu-Chek® Aviva glucometer (Roche, Basel, Switzerland) and the blood lactate concentration (mM) was analyzed using an enzyme electrode method with Lactate PRO 2 (Arkray Inc, Kyoto, Japan).

##### Training

All animals trained for 4 weeks, 5 days/week (Monday to Friday). Training protocols were adapted from our previous works ^13,14^ in terms of intensity and duration of sessions. To reduce animal anxiety, mice were trained in groups of four animals sharing the same cage.

Briefly, endurance training sessions started with identical warm-up as for the maximal endurance performance test. Then, all sessions of continuous running had a mean duration of 60 min and the distance covered every day was 1000 m, as a fixed volume. However, the intensity in terms of maximal speed, number of stages, as well as the speed and duration of each stage, varied along the week according to this structure: 2 days at high intensity (Tuesday and Friday), 2 days at moderate intensity (Monday and Thursday), and 1 day at low intensity (Wednesday). Speed ranged from 20 to 40 cm/s, which corresponded to 40-80% of mean maximal speed at the pre-training test ^54^. The duration of each stage varied inversely with speed, between 15 and 5 min ^54^. The slope was fixed at 10°.

Resistance training sessions started with an identical warm-up as for the maximal resistance performance test. Then, all sessions were designed to achieve the same exercise volume by means of a combination of number of steps climbed (or distance against gravity) and weight load ^55^. The number of steps per training session varied between 400-2000 depending on the maximal weight load, which ranged between 20-50 g or 25-65% of the maximal weight load at the pre-training test. We selected these maximum weight ranges because it has been described that below 75% of 1 repetition maximum there is no velocity loss, which is important for standardizing intensity of submaximal efforts ^56^. Week planning was: 2 days with high weight load and low number of steps (Tuesday and Friday), 2 days of intermediate weight load and number of steps (Monday and Thursday), and 1 day without weight load but a high number of steps (Wednesday). Control mice remained in a cage, in the same room, while END and RES mice were training.

#### Micro-computed tomography (MCT) scan

Micro-computed tomography (CT) scans were performed on unconscious mice using Argus PET/CT (Sedecal, Servicios Científico-Técnicos, University of Oviedo, Spain), to assess the muscle area of the hind leg of 1 CON, 1 miR-29KO, 1 KOEND and 1 KORES mouse. Prior to analysis mice were anesthetized with a continuous flow of a 3% isoflurane/oxygen mixture (IsoFlo, Zoetis España, Alcobendas, Madrid) and placed in prone position, leaving the hind limbs in a natural position. The analysis time was 10 minutes, and the slice thickness was 1.123 mm. The images were downloaded to the program RadiAnt DICOM Viewer [Software] (Version 2021.1). Muscle area was measured in a single axial cut placed in the middle part of the right leg. The specific anatomical reference was the middle point between the distal and proximal epiphyses of the tibia. Muscle area was assessed using the software Confocal Uniovi Fiji ImageJ (1.6 version), by manually drawing the muscle perimeter of lower leg and subtracting the area o tibial and fibular bones. The area was normalized to the body weight of the mouse.

#### MiR-29a-3p inhibitor transfection

To fine-tune the transfection protocol, 2 days after induction of differentiation, C2C12 cells were transfected with two different concentrations (15 nM or 100 nM) of mmu-miR-29a-3p MirVana inhibitors or a negative control inhibitor (Life Technologies, Carlsbad, CA, USA). Transfection at the same concentration was repeated at 48h. Each transfection was performed for 24 h in Opti-MEM™ I Reduced-Serum Medium (31985062, Gibco, Thermo Fisher Scientific, Inc., MA, USA) with Lipofectamine™ RNAiMAX Transfection Reagent (13778150, Invitrogen, Thermo Fisher Scientific, Inc., MA, USA). This is an improved Minimal Essential Medium (MEM) that allows for a reduction of FBS supplementation by at least 50% with no change in growth rate or morphology. The transfection efficiency of the different concentrations of miR-29a-3p inhibitors was estimated by qPCR, reflecting the inhibition activity of the different concentrations of miR-29a-3p inhibitors. Considering the transfection efficiency, the 100 nM concentration was selected for a second protocol in which, after transfection on days 2 and 4 of differentiation, cells were electrostimulated on day 6, following the same protocol described in electrical pulse stimulation section.

#### Electrical pulse stimulation (EPS)

On day 6 of differentiation, differentiated myotubes were electrically stimulated using the C-Pace EP cell culture stimulator with carbon electrodes ^16^ (EPS; C-Pace EP; Ionoptix, MA, USA). To simulate basal exercise conditions, 24 hours before electrostimulation, the culture media was changed to DMEM with low glucose concentration (1 g/L) and EVs-free HS to avoid contamination of EVs. Both electrostimulated and control cells remained in this media during the protocol. The EPS protocol consisted of applying pulses continuously (2 ms) to the myotubes at 1 Hz and 40V for 3 h in the incubator (at 37°C and 5% CO_2_). The contractions of the myotubes were observed under a microscope. Both the electrostimulation and control plates remained with the electrodes. However, in the case of the controls, there was no stimulation. Once the simulation period was completed, the medium samples were collected for EVs isolation, the plates were washed with PBS and cell lysis performed 0, 3 and 20 h after EPS.

#### Cell viability

The lactate dehydrogenase (LDH) release assay was used as a colorimetric method to estimate cell cytotoxicity. The presence of LDH in the extracellular media reveals a defect in cell membrane permeability due to cell death. Briefly, cells were exposed to EPS protocol for 3 hours under normal conditions and under conditions of miR-29a-3p inhibition. 100 μL of the supernatant from each of the conditions was transferred to a 96-well plate and 100 μL of LDH assay reagent consisting of a dye and a catalyser were added to each well. After an incubation period of 30 minutes under shaking and at room temperature, the absorbance was measured at 490 nm using microplate Versa Max Plate reader (MolecuarDevices, San José, California, United States). The absorbance at 690 nm was also measured as a reference for the background of each well.

As a positive control, 20 μL of a 0.1% Triton solution was added to the cells for 4 hours in the incubator at 37°C. Cells were then lysed and centrifuged at 500 x g for 5 minutes to remove cell debris. The supernatant was used as a positive control. Cell death was expressed as the percentage of cytotoxicity with respect to the positive control for each condition.

#### Sampling

##### Animals

All mice were sacrificed 24 h after the last training session. Mice were deeply anesthetized with ketamine (10 mg/mL) and xylazine (1 mg/mL) in saline solution and the different tissues were obtained for subsequent analysis. Blood was drawn from the vena cava of each mouse, using an intravascular catheter (0.9 x 25 mm) (BD Insyte, 381223) pretreated with EDTA (0.5 M) as an anticoagulant. After collection, plasma was rapidly separated by centrifugation at 2500 × g for 15 min at 4 °C. The plasma samples were immediately frozen in liquid nitrogen and stored at −80°C.

After blood sample collection, a cut was made in the abdominal aorta and all mice were perfused with 20 ml of cold PBS from the same blood collection site, the vena cava, to flush the tissues and prevent the blood interference present in the tissues in the ulterior miRNA analysis.

Skeletal muscles (quadriceps and soleus) and liver of each animal were removed. Right quadriceps and soleus were extracted, washed in PBS, cleaned of connective tissue, rapidly frozen in liquid nitrogen, and stored at −80°C for biochemical analysis.

##### Cell culture

From the culture media of each of the plates (approximately 12 mL), total EVs were isolated using total EV isolation reagent (exosome-enriched) (4478359, Thermo Fisher Scientific, Inc., MA, USA). The reagent was added to the total volume of culture media at a ratio of 1:2 and the solution was incubated overnight at 4°C. Precipitated exosomes were recovered by standard centrifugation at 10,000 x g for 60 min. An aliquot of the supernatant was taken as a negative control for the presence of EVs and the remainder was removed. The pellet was resuspended in 200 µl of modified PBS (HyClone Dulbecco’s phosphate buffered saline: liquid, Cytiva SH30028.02, Thermo Fisher Scientific, Inc., MA, USA), an aliquot was used to measure protein concentration by BSA technique, and the remainder was frozen at −80 °C until use.

After removal the culture media, cells were washed with PBS (1X) and used to extract RNA (2 × 10^6^ cells). 700 µl of Qiazol lysis reagent (QIAGEN, Hilden, Germany) was added to the cell culture plate. Cells were detached used a scraper and transfer to 1.5 mL tube. The content was shaken vigorously with the vortex for 1 minute for further analysis.

#### Sequencing and validation of mice EV plasma miRNA samples

100 μL of plasma from 5 samples per group (CON, END and RES) were pooled, generating three pools of 500 μL each. From this plasma volume (500 μL) EVs precipitation was conducted with exoRNeasy (QIAGEN, Hilden, Germany) after manufacturer’s instructions. RNA was isolated from these plasma EVs with proprietary RNA isolation protocol (Exiqon Services, Denmark) optimized for serum/plasma (no carrier added). Total RNA was eluted in ultra-low volume. The library preparation was done using the QIAseq miRNA Library Kit (QIAGEN, Hilden, Germany). A total of 5µl total RNA was converted into microRNA next generation sequencing (NGS) libraries. Adapters containing unique molecular identifiers (UMIs) were ligated to the RNA. Then RNA was converted to cDNA using the miRCURY LNA RT kit (QIAGEN, Hilden, Germany). The cDNA was amplified using PCR (22 cycles) and during the PCR indices were added. After PCR the samples were purified. Library preparation QC was performed using either Bioanalyzer 2100 (Agilent, Santa Clara, CA, USA) or TapeStation 4200 (Agilent). Based on quality of the inserts and the concentration measurements the libraries were pooled in equimolar ratios. The library pool(s) were quantified using the qPCR ExiSEQ LNA™ Quant kit (Exiqon, Denmark). The library pools were then sequenced on a NextSeq500 sequencing instrument according to the manufacturer instructions. Raw data was demultiplexed and FASTQ files for each sample were generated using the bcl2fastq conversion software v2.20 (Illumina inc., San Diego, CA, USA). FASTQ data were checked using the FastQC tool. Bowtie2 was used for sequence alignment against reference miRNA sequences from mice. Feature counting was carried out using HTSeq-count. Before differential expression analysis, the obtained count matrix was normalized using the TMM method available within the edgeR library from the R coding environment. Raw data are available on ZENODO. (https://zenodo.org/doi/10.5281/zenodo.11184908).

#### Total RNA extraction

##### Plasma EVs

A selection of miRNAs was carried out from NGS data with the criteria of highly expressed miRNAs with a moderate to high relative expression change difference regarding CON, more than 1000 RPM and a log2 FC higher than 0.5.

Secondly, the volume not pooled of every single plasma sample from CON, END and RES was treated with thrombin to create serum like samples (SBI, Palo Alto, CA, USA). SeraMiR Exosome RNA purification kit (SBI, Palo Alto, CA, USA) was used to facilitated extracellular vesicles (EVs) precipitation and extraction. miRCURY RNA isolation kit (QIAGEN Hilden, Germany) was used to extract miRNAs from 200 µl of plasma samples following the manufacturer’s instructions. The RNA Spike-in kit (QIAGEN Hilden, Germany) was used, which contains several synthetic RNA spike-in templates. This kit includes *Caenorhabditis elegans* synthetic miR-39-3p (cel-miR-39-3p), which lacks sequence homology with mouse miRNAs and was used for external normalization. In addition, it also contains synthetic RNA spike-in templates (UniSp2, UniSp4, UniSp5), which were used in all extractions to control the efficiency of RNA isolation. RNA was eluted in 30 μl RNase-free H_2_O and its concentration in the samples were measured by NanoDrop (Thermo Fisher Scientific, Inc., MA, USA), then stored in a −80°C freezer.

##### Cell culture EVs

In the case of EVs obtained from the culture media, RNA extraction was performed using the Total Exosome RNA & Protein Isolation Kit (4478545, Invitrogen, Thermo Fisher Scientific, Inc., MA, USA). First the 200 µL aliquot of EVs was added a 2X volume of Denaturing Solution and mixed. The mixture was incubated on ice for 5 minutes. A volume of phenol-chloroform was added, and the samples were vortexed for 30 seconds. To separate the aqueous phase (RNA) from the organic phase, samples were centrifuged for 5 minutes at maximum speed (> 10,000 x g) at room temperature. The aqueous phase was carefully collected, and 1.25 volumes of absolute ethanol were added to the aqueous phase. The total sample was transferred to a column and centrifuged at 10,000 x g for 12 seconds. This was followed by a series of washes in which the centrifugation conditions were the same. RNA was eluted in 50 µL in preheated RNase-free H_2_O. RNA concentration was measured by Nanodrop and stored at −80°C.

##### Animal tissues and C2C12 cells

In dry ice conditions, 50 mg of quadriceps, 25 mg of liver and 5 mg of soleus were cut, and 2 x 10^6^ C2C12 cells were collected to perform RNA extraction. Tissues were disrupted using a mortar and pestle with 700 µl of Qiazol lysis reagent (QIAGEN, Hilden, Germany). MiRNeasy mini kit (QIAGEN, Hilden, Germany) for quadriceps and liver and cell pellet and miRNeasy micro kit (QIAGEN, Hilden, Germany) for soleus were used to extract total RNA following the manufacturer’s instructions. The RNA Spike-in kit with synthetic RNA spike-in templates (QIAGEN, Hilden, Germany) was added in all extractions to monitor RNA isolation efficiency. RNA was eluted in 30 µl RNase-free H_2_O. All samples were measured by NanoDrop (Thermo Fisher Scientific, Inc., MA, USA) and stored at −80°C to further analysis.

#### RT-qPCR for miRNAs

cDNA was synthesized using the miRCURY LNA RT kit (QIAGEN, Hilden, Germany) ^57^. For plasma EVs, 2 µl RNA samples were reverse transcribed in 10 µl reaction. For tissue, EVs and C2C12 cells analysis equal quantities of RNA (5 ng/µL) were used. Additional spike-in (UniSp6) (QIAGEN, Hilden, Germany) was added to the cDNA synthesis reaction to check for RT and PCR inhibitors. RT reaction was performed with the following conditions: incubation for 60 min at 42°C, heat-inactivation for 5 min at 95°C, immediately cool to 4°C. For qPCR, cDNA was diluted 1:40 and 4 µl used in 10 µl with miRCURY LNA SYBR Green PCR master mix (QIAGEN, Hilden, Germany) on a 7900HT fast real-time PCR system (Applied Biosystems, Waltham, MA, USA) with the following cycling conditions: 10 min at 95 °C, 40 cycles of 10 s at 95 °C and 1 min at 60 °C, followed by melting curve analysis. Primers for miRNAs were obtained from Qiagen (Table S1). UniSp6 was used as an internal reference for miRNA expression. Relative levels of miRNAs and were quantitatively analyzed using the 2^−ΔΔCt^method ^58^. The mean expression of all miRNAs analyzed for each sample was used as an internal experimental control.

#### RT-qPCR for mRNAs

cDNA was synthesised using the StaRT Reverse Transcription kit (AnyGenes, Paris, France) for mRNA. For tissue analysis equal quantities of RNA (500 ng) were used. For mRNA, cDNA was diluted 6-fold and 1 µL was used in 10 µL qPCR reactions with the Perfect Master Mix SYBR Green kit (AnyGenes, Paris, France) in the same 7900HT real-time PCR system (Applied Biosystems, Waltham, MA, USA) with the following cycling conditions: 10 min at 95 °C, 40 cycles of 10 s at 95 °C and 30 s at 60 °C and 1 min at 60 °C, and 30 s at 65 °C. Relative levels of genes were quantitatively analyzed using the 2^−ΔΔCt^ method^58^. Primers for the genes analyzed were specifically designed by AnyGenes; the information is listed in STAR Methods table. β*-actin* and *Gapdh* were used as housekeeping gene for mRNA expression.

#### Western Blot

For tissue samples, 20 mg of quadriceps and 5 mg of soleus were homogenized in 400 μL and 100 μL of homogenization buffer respectively, consisting of: 20 mM HEPES, pH 7.4, 100 mM NaCl, 50 mM NaF, 5 mM EDTA, 1% Triton X-100, 1 mM sodium orthovanadate, 1 mM pyrophosphate and 1X Complete™ Protease Inhibitor Cocktail (CO-RO Roche Life Science, Basel, Switzerland). Samples were then centrifuged at 12,000 x g at 4°C for 10 min and the supernatant was collected and stored at −80°C until use. For the specific analysis of mitochondrial complexes, three freeze/thaw cycles were performed after homogenization to promote the rupture not only of the plasma membrane but also the mitochondrial membrane: 1) 1h at −80°C and thawing at 4°C; 2) 24h at −80°C and thawing at 4°C; and 3) 1 h at 80°C and thawing at 4°C.

For EVs samples, 200 μL of sample containing EVs dissolved in PBS 1X were used. Homogenization was performed by sonication with continuous 35% pulses of 10 seconds on and 5 seconds off for 1 minute.

Protein quantification was determined by Nanodrop. Total protein (between 30-50 μg of tissue and 10 μg of cell pellet and EVs) was resuspended in loading buffer (BR5X) (300 mM Tris/HCl, pH 6.8, 10% SDS, 50% glycerol, 20% 2-mercaptoethanol, 40 mM EDTA and 0.05% bromophenol). Prior to gel loading, samples were boiled in a thermoblock for 5 min at 95°C (except those sample used for mitochondrial complexes analysis) and loaded onto 4-12% precast Bis-Tris gels (MP41G12, MerckMillipore, Darmstadt, Germany). In addition to the samples, the Prestained Protein Ladder marker with a range of 5-245 kDa (AB116029, Abcam, Cambridge, UK) was also loaded onto the gels. Proteins were transferred to PVDF membranes (pore size of 0.2 µm, 15239814, Amersham Hybond^TM^, Amersham, UK). Transfer was carried out wet with the transfer buffer at a constant amperage of 250 mA at 4°C. After transfer, the membranes were incubated in blocking solution (5% bovine serum albumin, BSA) in TBS-T (200 mM Tris/HCl, pH 7.6, 1.5M NaCl and Tween 20) at room temperature and shaking. After blocking, membranes were incubated with the corresponding primary antibody (see table STAR Methods) in TBS-T 1X overnight at 4°C and shaking. Next day membranes were washed with TBS-T 1X and incubated with the appropriate horseradish peroxidase-conjugated secondary antibodies (diluted 1:20,000) for 1 hour at room temperature. Quantification of relative protein levels was performed by reflecting this level as a ratio of each protein of interest to the housekeeping protein using the protocol developed by Hossein Davarineja^59^ in Uniovi Fiji Confocal software (coupled to ImageJ, version 1.6).

#### In silico

Pathway analysis of the genes targeted by the EV miRNAs identified was performed to gain insight into their potential functional implication in the biological response to exercise. For each miRNA, experimentally validated targets were retrieved from miRTarBase and miRWalk databases. TarBase v7.0 validated gene targets were used on DIANA TOOLS miRpath v.3 using KEGG pathways union analysis^60,61^. We used target mining analysis by miRWalk for gene interactions. The data downloaded from miRWalk were utilized on Cytoscape v3.10.0 to create target intersection between miRNAs.

#### Statistical Analysis

Unless otherwise specified, data are presented as mean ± SEM and represented as column scatter plots. Normality of the variables was tested by means of the Shapiro[Wilk test. Thus, for the variables that met these assumptions, Student’s t-test for independent or paired samples and analysis of variance (ANOVA), followed by a Tukey post hoc analysis, were applied, depending on the number of groups to be compared. For variables that did not meet the assumptions of normality, Mann-Whitney U or Kruskal-Wallis tests were compared followed by a Dunn post hoc analysis, depending on the number of groups to be compared. Pearson’s correlation coefficient was calculated to explore associations between variables. p ≤ 0.05 was considered significant, and statistically significant differences are shown in each graph with the p value. All statistical analyses and graphical representations were carried out with SPSS software version 27 for Windows (IBM, Armonk, NY, USA) and GraphPad Prism version 8.0.1 for Windows (GraphPad, San Diego, CA, USA), respectively. The correlation coefficient plots were generated using the SRPLOT online tool^62^.

#### Data and code availability

Raw RNA-Seq data have been deposited at Zenodo and are publicly accessible as of the date of publication. Accession link is listed in the key resources table.

All the values used to create graphs in the paper can be found in the Data Main Figures file in a single Excel spreadsheet.

The raw uncropped western blots can be found in the Data Supplemental Figures file in a single PDF document.

Additional information required to reanalyze the data reported in this paper is available from the lead contact upon request.

## Notes

https://zenodo.org/doi/10.5281/zenodo.11184908

### Competing Interest Statement

The authors have declared no competing interest.

## REFERENCES

1. Thompson, W.R., Sallis, R., Joy, E., Jaworski, C.A., Stuhr, R.M., and Trilk, J.L. (2020). Exercise Is Medicine. Am J Lifestyle Med 14, 511–523. 10.1177/1559827620912192.

2. Safdar, A., Saleem, A., and Tarnopolsky, M.A. (2016). The potential of endurance exercise-derived exosomes to treat metabolic diseases. Nat Rev Endocrinol 12, 504–517. 10.1038/nrendo.2016.76.

3. Brahmer, A., Neuberger, E., Esch-Heisser, L., Haller, N., Jorgensen, M.M., Baek, R., Mobius, W., Simon, P., and Kramer-Albers, E.M. (2019). Platelets, endothelial cells and leukocytes contribute to the exercise-triggered release of extracellular vesicles into the circulation. J Extracell Vesicles 8, 1615820. 10.1080/20013078.2019.1615820.

4. Whitham, M., Parker, B.L., Friedrichsen, M., Hingst, J.R., Hjorth, M., Hughes, W.E., Egan, C.L., Cron, L., Watt, K.I., Kuchel, R.P., et al. (2018). Extracellular Vesicles Provide a Means for Tissue Crosstalk during Exercise. Cell metabolism 27, 237–251 e234. 10.1016/j.cmet.2017.12.001.

5. Abels, E.R., and Breakefield, X.O. (2016). Introduction to Extracellular Vesicles: Biogenesis, RNA Cargo Selection, Content, Release, and Uptake. Cell Mol Neurobiol 36, 301–312. 10.1007/s10571-016-0366-z.

6. O’Brien, K., Breyne, K., Ughetto, S., Laurent, L.C., and Breakefield, X.O. (2020). RNA delivery by extracellular vesicles in mammalian cells and its applications. Nature reviews 21, 585–606. 10.1038/s41580-020-0251-y.

7. Estebanez, B., Jimenez-Pavon, D., Huang, C.J., Cuevas, M.J., and Gonzalez-Gallego, J. (2021). Effects of exercise on exosome release and cargo in in vivo and ex vivo models: A systematic review. J Cell Physiol 236, 3336–3353. 10.1002/jcp.30094.

8. Darragh, I.A.J., O’Driscoll, L., and Egan, B. (2021). Exercise Training and Circulating Small Extracellular Vesicles: Appraisal of Methodological Approaches and Current Knowledge. Front Physiol 12, 738333. 10.3389/fphys.2021.738333.

9. Isaac, R., Reis, F.C.G., Ying, W., and Olefsky, J.M. (2021). Exosomes as mediators of intercellular crosstalk in metabolism. Cell metabolism 33, 1744–1762. 10.1016/j.cmet.2021.08.006.

10. Ebert, M.S., and Sharp, P.A. (2012). Roles for microRNAs in conferring robustness to biological processes. Cell 149, 515–524. 10.1016/j.cell.2012.04.005.

11. Yanez-Mo, M., Siljander, P.R., Andreu, Z., Zavec, A.B., Borras, F.E., Buzas, E.I., Buzas, K., Casal, E., Cappello, F., Carvalho, J., et al. (2015). Biological properties of extracellular vesicles and their physiological functions. J Extracell Vesicles 4, 27066. 10.3402/jev.v4.27066.

12. Murphy, R.M., Watt, M.J., and Febbraio, M.A. (2020). Metabolic communication during exercise. Nat Metab 2, 805–816. 10.1038/s42255-020-0258-x.

13. Fernandez, J., Fernandez-Sanjurjo, M., Iglesias-Gutierrez, E., Martinez-Camblor, P., Villar, C.J., Tomas-Zapico, C., Fernandez-Garcia, B., and Lombo, F. (2021). Resistance and Endurance Exercise Training Induce Differential Changes in Gut Microbiota Composition in Murine Models. Front Physiol 12, 748854. 10.3389/fphys.2021.748854.

14. Codina-Martinez, H., Fernandez-Garcia, B., Diez-Planelles, C., Fernandez, A.F., Higarza, S.G., Fernandez-Sanjurjo, M., Diez-Robles, S., Iglesias-Gutierrez, E., and Tomas-Zapico, C. (2020). Autophagy is required for performance adaptive response to resistance training and exercise-induced adult neurogenesis. Scand J Med Sci Sports 30, 238–253. 10.1111/sms.13586.

15. Castano, C., Mirasierra, M., Vallejo, M., Novials, A., and Parrizas, M. (2020). Delivery of muscle-derived exosomal miRNAs induced by HIIT improves insulin sensitivity through down-regulation of hepatic FoxO1 in mice. Proceedings of the National Academy of Sciences of the United States of America 117, 30335–30343. 10.1073/pnas.2016112117.

16. Abdelmoez, A.M., Sardon Puig, L., Smith, J.A.B., Gabriel, B.M., Savikj, M., Dollet, L., Chibalin, A.V., Krook, A., Zierath, J.R., and Pillon, N.J. (2020). Comparative profiling of skeletal muscle models reveals heterogeneity of transcriptome and metabolism. Am J Physiol Cell Physiol 318, C615–C626. 10.1152/ajpcell.00540.2019.

17. Dastah, S., Tofighi, A., and Bonab, S.B. (2021). The effect of aerobic exercise on the expression of mir-126 and related target genes in the endothelial tissue of the cardiac muscle of diabetic rats. Microvasc Res 138, 104212. 10.1016/j.mvr.2021.104212.

18. Rubenstein, A.B., Smith, G.R., Raue, U., Begue, G., Minchev, K., Ruf-Zamojski, F., Nair, V.D., Wang, X., Zhou, L., Zaslavsky, E., et al. (2020). Single-cell transcriptional profiles in human skeletal muscle. Sci Rep 10, 229. 10.1038/s41598-019-57110-6.

19. Kavakiotis, I., Alexiou, A., Tastsoglou, S., Vlachos, I.S., and Hatzigeorgiou, A.G. (2022). DIANA-miTED: a microRNA tissue expression database. Nucleic Acids Res 50, D1055–D1061. 10.1093/nar/gkab733.

20. Caravia, X.M., Fanjul, V., Oliver, E., Roiz-Valle, D., Moran-Alvarez, A., Desdin-Mico, G., Mittelbrunn, M., Cabo, R., Vega, J.A., Rodriguez, F., et al. (2018). The microRNA-29/PGC1alpha regulatory axis is critical for metabolic control of cardiac function. PLoS Biol 16, e2006247. 10.1371/journal.pbio.2006247.

21. Massart, J., Sjogren, R.J.O., Lundell, L.S., Mudry, J.M., Franck, N., O’Gorman, D.J., Egan, B., Zierath, J.R., and Krook, A. (2017). Altered miR-29 Expression in Type 2 Diabetes Influences Glucose and Lipid Metabolism in Skeletal Muscle. Diabetes 66, 1807–1818. 10.2337/db17-0141.

22. Kriegel, A.J., Liu, Y., Fang, Y., Ding, X., and Liang, M. (2012). The miR-29 family: genomics, cell biology, and relevance to renal and cardiovascular injury. Physiol Genomics 44, 237–244. 10.1152/physiolgenomics.00141.2011.

23. Liu, C., Li, L., Ge, M., Gu, L., Zhang, K., Su, Y., Zhang, Y., Liu, C., Lan, M., Yu, Y., et al. (2021). MiR-29ab1 Cluster Resists Muscle Atrophy Through Inhibiting MuRF1. DNA Cell Biol 40, 1167–1176. 10.1089/dna.2021.0267.

24. Ferreira, J.C., Rolim, N.P., Bartholomeu, J.B., Gobatto, C.A., Kokubun, E., and Brum, P.C. (2007). Maximal lactate steady state in running mice: effect of exercise training. Clin Exp Pharmacol Physiol 34, 760–765. 10.1111/j.1440-1681.2007.04635.x.

25. Willows, R., Sanders, M.J., Xiao, B., Patel, B.R., Martin, S.R., Read, J., Wilson, J.R., Hubbard, J., Gamblin, S.J., and Carling, D. (2017). Phosphorylation of AMPK by upstream kinases is required for activity in mammalian cells. The Biochemical journal 474, 3059–3073. 10.1042/BCJ20170458.

26. Richter, E.A., and Hargreaves, M. (2013). Exercise, GLUT4, and skeletal muscle glucose uptake. Physiol Rev 93, 993–1017. 10.1152/physrev.00038.2012.

27. Guo, L., Quan, M., Pang, W., Yin, Y., and Li, F. (2023). Cytokines and exosomal miRNAs in skeletal muscle-adipose crosstalk. Trends Endocrinol Metab 34, 666–681. 10.1016/j.tem.2023.07.006.

28. Ni, P., Yang, L., and Li, F. (2023). Exercise-derived skeletal myogenic exosomes as mediators of intercellular crosstalk: a major player in health, disease, and exercise. J Physiol Biochem 79, 501–510. 10.1007/s13105-023-00969-x.

29. Fischetti, F., Poli, L., De Tommaso, M., Paolicelli, D., Greco, G., and Cataldi, S. (2023). The role of exercise parameters on small extracellular vesicles and microRNAs cargo in preventing neurodegenerative diseases. Front Physiol 14, 1241010. 10.3389/fphys.2023.1241010.

30. Darragh, I.A.J., and Egan, B. (2024). Considerations for exerkine research focusing on the response to exercise training. J Sport Health Sci 13, 130–132. 10.1016/j.jshs.2023.11.002.

31. Watanabe, S., Sudo, Y., Makino, T., Kimura, S., Tomita, K., Noguchi, M., Sakurai, H., Shimizu, M., Takahashi, Y., Sato, R., and Yamauchi, Y. (2022). Skeletal muscle releases extracellular vesicles with distinct protein and microRNA signatures that function in the muscle microenvironment. PNAS Nexus 1, pgac173. 10.1093/pnasnexus/pgac173.

32. Lautaoja-Kivipelto, J.H., Karvinen, S., Korhonen, T.M., O’Connell, T.M., Tiirola, M., Hulmi, J.J., and Pekkala, S. (2024). Interaction of the C2C12 myotube contractions and glucose availability on transcriptome and extracellular vesicle microRNAs. Am J Physiol Cell Physiol 326, C348–C361. 10.1152/ajpcell.00401.2023.

33. Vann, C.G., Zhang, X., Khodabukus, A., Orenduff, M.C., Chen, Y.H., Corcoran, D.L., Truskey, G.A., Bursac, N., and Kraus, V.B. (2022). Differential microRNA profiles of intramuscular and secreted extracellular vesicles in human tissue-engineered muscle. Front Physiol 13, 937899. 10.3389/fphys.2022.937899.

34. Soriano-Arroquia, A., McCormick, R., Molloy, A.P., McArdle, A., and Goljanek-Whysall, K. (2016). Age-related changes in miR-143-3p:Igfbp5 interactions affect muscle regeneration. Aging Cell 15, 361–369. 10.1111/acel.12442.

35. Wei, W., He, H.B., Zhang, W.Y., Zhang, H.X., Bai, J.B., Liu, H.Z., Cao, J.H., Chang, K.C., Li, X.Y., and Zhao, S.H. (2013). miR-29 targets Akt3 to reduce proliferation and facilitate differentiation of myoblasts in skeletal muscle development. Cell Death Dis 4, e668. 10.1038/cddis.2013.184.

36. Guess, M.G., Barthel, K.K., Harrison, B.C., and Leinwand, L.A. (2015). miR-30 family microRNAs regulate myogenic differentiation and provide negative feedback on the microRNA pathway. PLoS ONE 10, e0118229. 10.1371/journal.pone.0118229.

37. Fabbri, M., Garzon, R., Cimmino, A., Liu, Z., Zanesi, N., Callegari, E., Liu, S., Alder, H., Costinean, S., Fernandez-Cymering, C., et al. (2007). MicroRNA-29 family reverts aberrant methylation in lung cancer by targeting DNA methyltransferases 3A and 3B. Proceedings of the National Academy of Sciences of the United States of America 104, 15805–15810. 10.1073/pnas.0707628104.

38. Rowlands, D.S., Page, R.A., Sukala, W.R., Giri, M., Ghimbovschi, S.D., Hayat, I., Cheema, B.S., Lys, I., Leikis, M., Sheard, P.W., et al. (2014). Multi-omic integrated networks connect DNA methylation and miRNA with skeletal muscle plasticity to chronic exercise in Type 2 diabetic obesity. Physiol Genomics 46, 747–765. 10.1152/physiolgenomics.00024.2014.

39. Schiaffino, S., and Reggiani, C. (2011). Fiber types in mammalian skeletal muscles. Physiol Rev 91, 1447–1531. 10.1152/physrev.00031.2010.

40. Nikolic, N., Bakke, S.S., Kase, E.T., Rudberg, I., Flo Halle, I., Rustan, A.C., Thoresen, G.H., and Aas, V. (2012). Electrical pulse stimulation of cultured human skeletal muscle cells as an in vitro model of exercise. PLoS ONE 7, e33203. 10.1371/journal.pone.0033203.

41. Valero-Breton, M., Warnier, G., Castro-Sepulveda, M., Deldicque, L., and Zbinden-Foncea, H. (2020). Acute and Chronic Effects of High Frequency Electric Pulse Stimulation on the Akt/mTOR Pathway in Human Primary Myotubes. Front Bioeng Biotechnol 8, 565679. 10.3389/fbioe.2020.565679.

42. Nikolic, N., Gorgens, S.W., Thoresen, G.H., Aas, V., Eckel, J., and Eckardt, K. (2017). Electrical pulse stimulation of cultured skeletal muscle cells as a model for in vitro exercise - possibilities and limitations. Acta Physiol (Oxf) 220, 310–331. 10.1111/apha.12830.

43. Hou, Z., Qin, X., Hu, Y., Zhang, X., Li, G., Wu, J., Li, J., Sha, J., Chen, J., Xia, J., et al. (2019). Longterm Exercise-Derived Exosomal miR-342-5p: A Novel Exerkine for Cardioprotection. Circ Res 124, 1386–1400. 10.1161/CIRCRESAHA.118.314635.

44. de Gonzalo-Calvo, D., Davalos, A., Fernandez-Sanjurjo, M., Amado-Rodriguez, L., Diaz-Coto, S., Tomas-Zapico, C., Montero, A., Garcia-Gonzalez, A., Llorente-Cortes, V., Heras, M.E., et al. (2018). Circulating microRNAs as emerging cardiac biomarkers responsive to acute exercise. Int J Cardiol 264, 130–136. 10.1016/j.ijcard.2018.02.092.

45. Davidsen, P.K., Gallagher, I.J., Hartman, J.W., Tarnopolsky, M.A., Dela, F., Helge, J.W., Timmons, J.A., and Phillips, S.M. (2011). High responders to resistance exercise training demonstrate differential regulation of skeletal muscle microRNA expression. J Appl Physiol (1985) 110, 309–317. 10.1152/japplphysiol.00901.2010.

46. Song, H., Ding, L., Zhang, S., and Wang, W. (2018). MiR-29 family members interact with SPARC to regulate glucose metabolism. Biochemical and biophysical research communications 497, 667–674. 10.1016/j.bbrc.2018.02.129.

47. Esteves, J.V., Enguita, F.J., and Machado, U.F. (2017). MicroRNAs-Mediated Regulation of Skeletal Muscle GLUT4 Expression and Translocation in Insulin Resistance. J Diabetes Res 2017, 7267910. 10.1155/2017/7267910.

48. Dos Santos, M., Backer, S., Aurade, F., Wong, M.M., Wurmser, M., Pierre, R., Langa, F., Do Cruzeiro, M., Schmitt, A., Concordet, J.P., et al. (2022). A fast Myosin super enhancer dictates muscle fiber phenotype through competitive interactions with Myosin genes. Nat Commun 13, 1039. 10.1038/s41467-022-28666-1.

49. Dreyer, H.C., Fujita, S., Cadenas, J.G., Chinkes, D.L., Volpi, E., and Rasmussen, B.B. (2006). Resistance exercise increases AMPK activity and reduces 4E-BP1 phosphorylation and protein synthesis in human skeletal muscle. J Physiol 576, 613–624. 10.1113/jphysiol.2006.113175.

50. Horita, M., Farquharson, C., and Stephen, L.A. (2021). The role of miR-29 family in disease. J Cell Biochem 122, 696–715. 10.1002/jcb.29896.

51. Roshan, R., Shridhar, S., Sarangdhar, M.A., Banik, A., Chawla, M., Garg, M., Singh, V.P., and Pillai, B. (2014). Brain-specific knockdown of miR-29 results in neuronal cell death and ataxia in mice. RNA 20, 1287–1297. 10.1261/rna.044008.113.

52. Hebert, S.S., Horre, K., Nicolai, L., Papadopoulou, A.S., Mandemakers, W., Silahtaroglu, A.N., Kauppinen, S., Delacourte, A., and De Strooper, B. (2008). Loss of microRNA cluster miR-29a/b-1 in sporadic Alzheimer’s disease correlates with increased BACE1/beta-secretase expression. Proceedings of the National Academy of Sciences of the United States of America 105, 6415–6420. 10.1073/pnas.0710263105.

53. Iglesias-Gutierrez, E., Fernandez-Sanjurjo, M., Fernandez, A.F., Rodriguez Diaz, F.J., Lopez-Taboada, I., Tomas-Zapico, C., and Fernandez-Garcia, B. (2021). Versatility of Protocols for Resistance Training and Assessment using Static and Dynamic Ladders in Animal Models. J Vis Exp. 10.3791/63098.

54. Kemi, O.J., Loennechen, J.P., Wisloff, U., and Ellingsen, O. (2002). Intensity-controlled treadmill running in mice: cardiac and skeletal muscle hypertrophy. J Appl Physiol (1985) 93, 1301–1309. 10.1152/japplphysiol.00231.2002.

55. Figueiredo, V.C., de Salles, B.F., and Trajano, G.S. (2018). Volume for Muscle Hypertrophy and Health Outcomes: The Most Effective Variable in Resistance Training. Sports Med 48, 499–505. 10.1007/s40279-017-0793-0.

56. Gentil, P., Marques, V.A., Neto, J.P.P., Santos, A.C.G., Steele, J., Fisher, J., Paoli, A., and Bottaro, M. (2018). Using velocity loss for monitoring resistance training effort in a real-world setting. Appl Physiol Nutr Metab 43, 833–837. 10.1139/apnm-2018-0011.

57. Mestdagh, P., Hartmann, N., Baeriswyl, L., Andreasen, D., Bernard, N., Chen, C., Cheo, D., D’Andrade, P., DeMayo, M., Dennis, L., et al. (2014). Evaluation of quantitative miRNA expression platforms in the microRNA quality control (miRQC) study. Nat Methods 11, 809–815. 10.1038/nmeth.3014.

58. Livak, K.J., and Schmittgen, T.D. (2001). Analysis of relative gene expression data using real-time quantitative PCR and the 2(-Delta Delta C(T)) Method. Methods 25, 402–408. 10.1006/meth.2001.1262.

59. Davarinejad, H. Quantifications of Western Blots with ImageJ. http://www.yorku.ca/yisheng/Internal/Protocols/ImageJ.pdf.

60. Vlachos, I.S., Paraskevopoulou, M.D., Karagkouni, D., Georgakilas, G., Vergoulis, T., Kanellos, I., Anastasopoulos, I.L., Maniou, S., Karathanou, K., Kalfakakou, D., et al. (2015). DIANA-TarBase v7.0: indexing more than half a million experimentally supported miRNA:mRNA interactions. Nucleic Acids Res 43, D153–159. 10.1093/nar/gku1215.

61. Vlachos, I.S., Zagganas, K., Paraskevopoulou, M.D., Georgakilas, G., Karagkouni, D., Vergoulis, T., Dalamagas, T., and Hatzigeorgiou, A.G. (2015). DIANA-miRPath v3.0: deciphering microRNA function with experimental support. Nucleic Acids Res 43, W460–466. 10.1093/nar/gkv403.

62. Tang, D., Chen, M., Huang, X., Zhang, G., Zeng, L., Zhang, G., Wu, S., and Wang, Y. (2023). SRplot: A free online platform for data visualization and graphing. PLoS ONE 18, e0294236. 10.1371/journal.pone.0294236.

