## Supplemental Figures for "miR-29a-3p, a new myokine orchestrating resistance exercise via coordinated metabolic responses"

Western blot analysis of plasma EVs

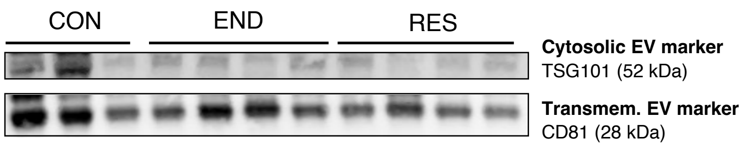


**Supplementary figure S1. Western blot analysis of the cytosolic EV marker TSG101 and the transmembrane EV marker CD81 EVs isolated from plasma samples of CON (n = 3), END (n = 4) and RES (n = 4) mice.**

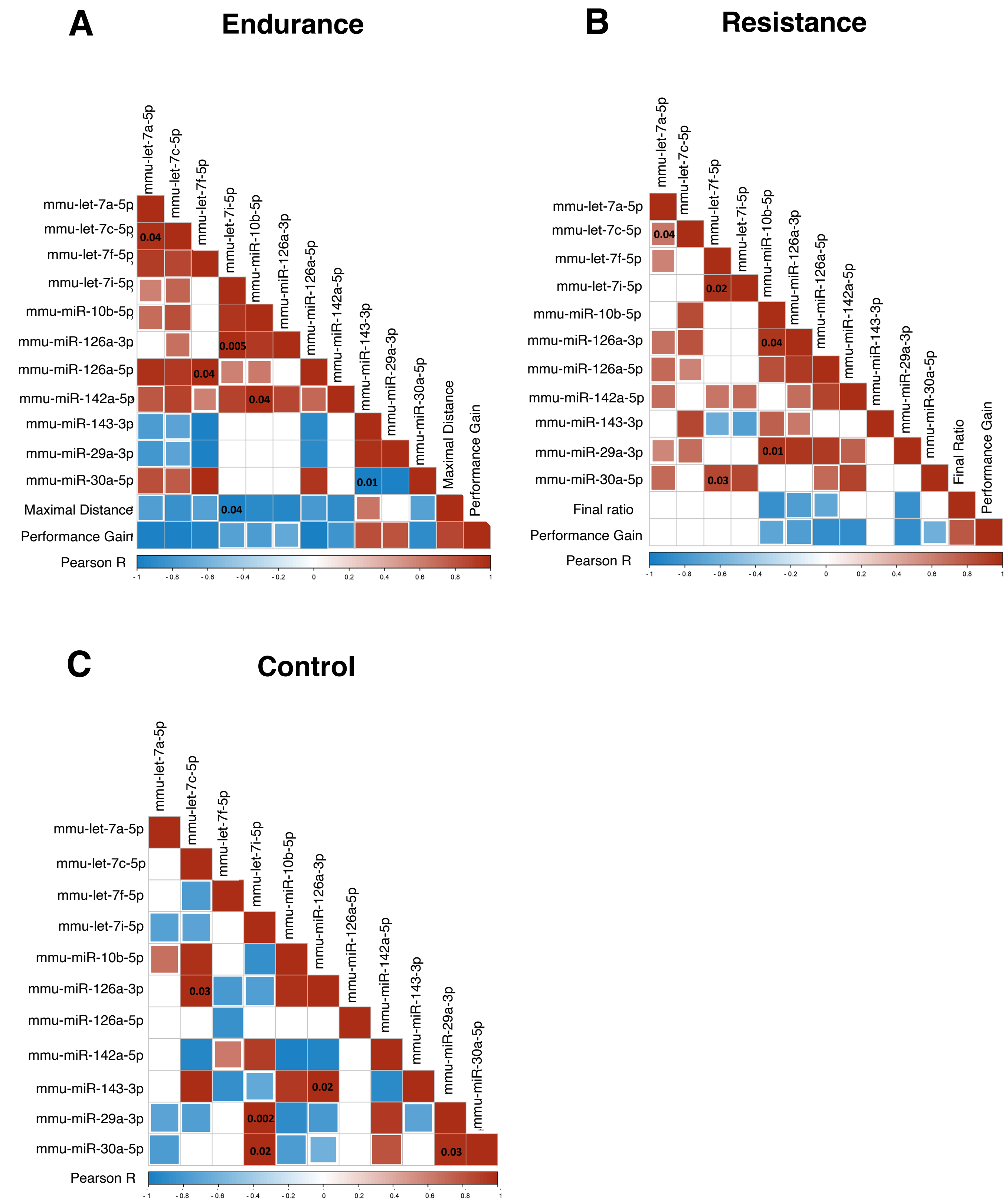


**Supplementary figure S2. Pearson correlation matrix between the expression levels of the 11 training-associated miRNAs in EVs in trained mice. (A)** END (n = 4). **(B)** RES (n = 5). **(C)** CON (n = 4). Blue boxes show a negative correlation (Pearson coefficient ≤ 1), and red boxes show a positive correlation (Pearson coefficient ≥ 1). Darker colors represent a stronger correlation and white colors indicate no correlation. The p-value is shown in the boxes.

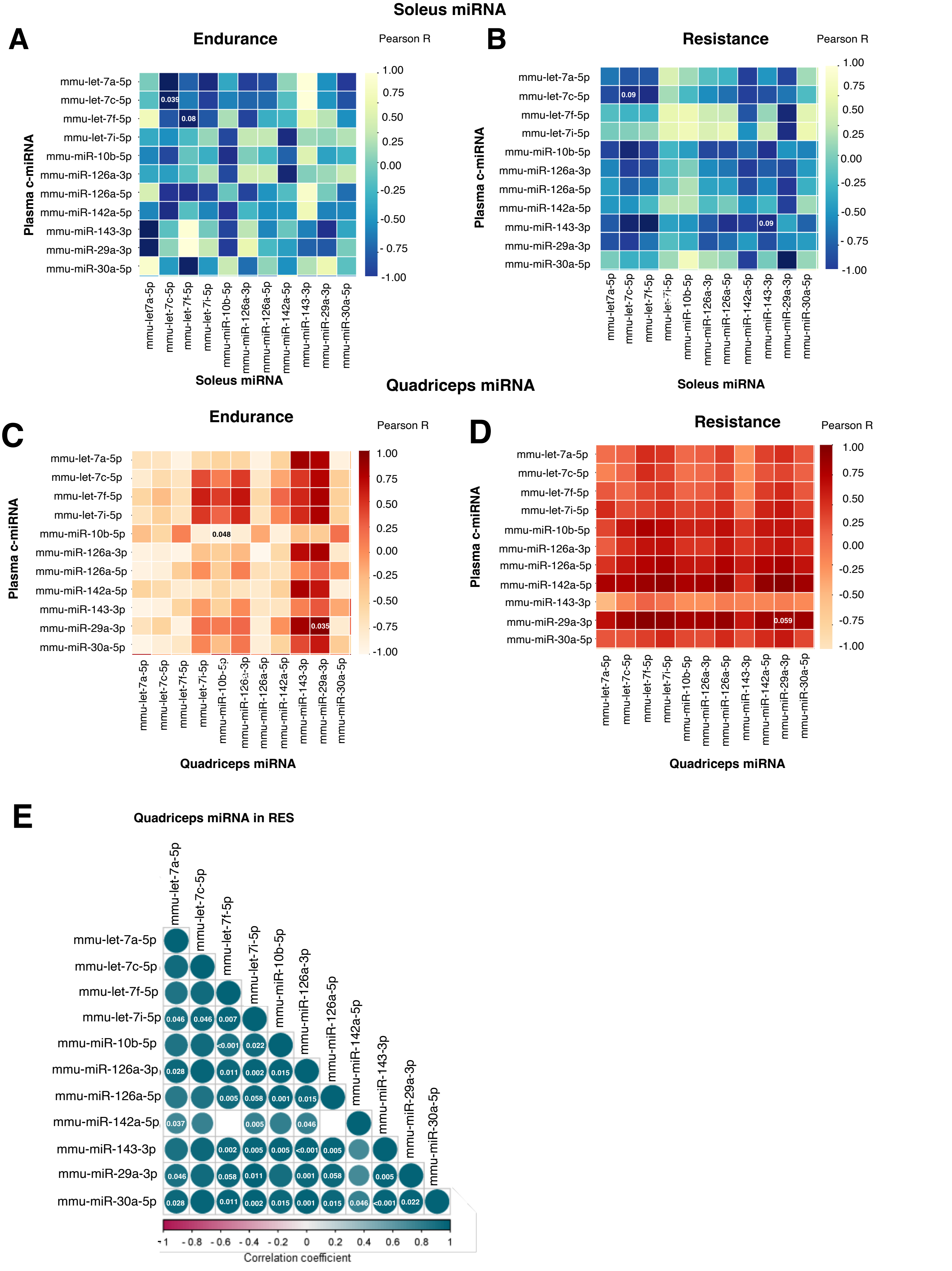


**Supplementary figure S3. Pearson correlation matrix between the expression levels of the 11 training-associated miRNAs in the soleus and in the quadriceps versus their levels in EVs and their levels in quadriceps of WT trained mice. (A)** Soleus END (n = 4). **(B)** Soleus RES (n = 5). **(C)** Quadriceps END (n = 4). **(D)** Quadriceps RES (n = 5). **(E)** Quadriceps RES (n = 8). The Pearson correlation is shown in the figures and the p-value is shown in the boxes and circles.

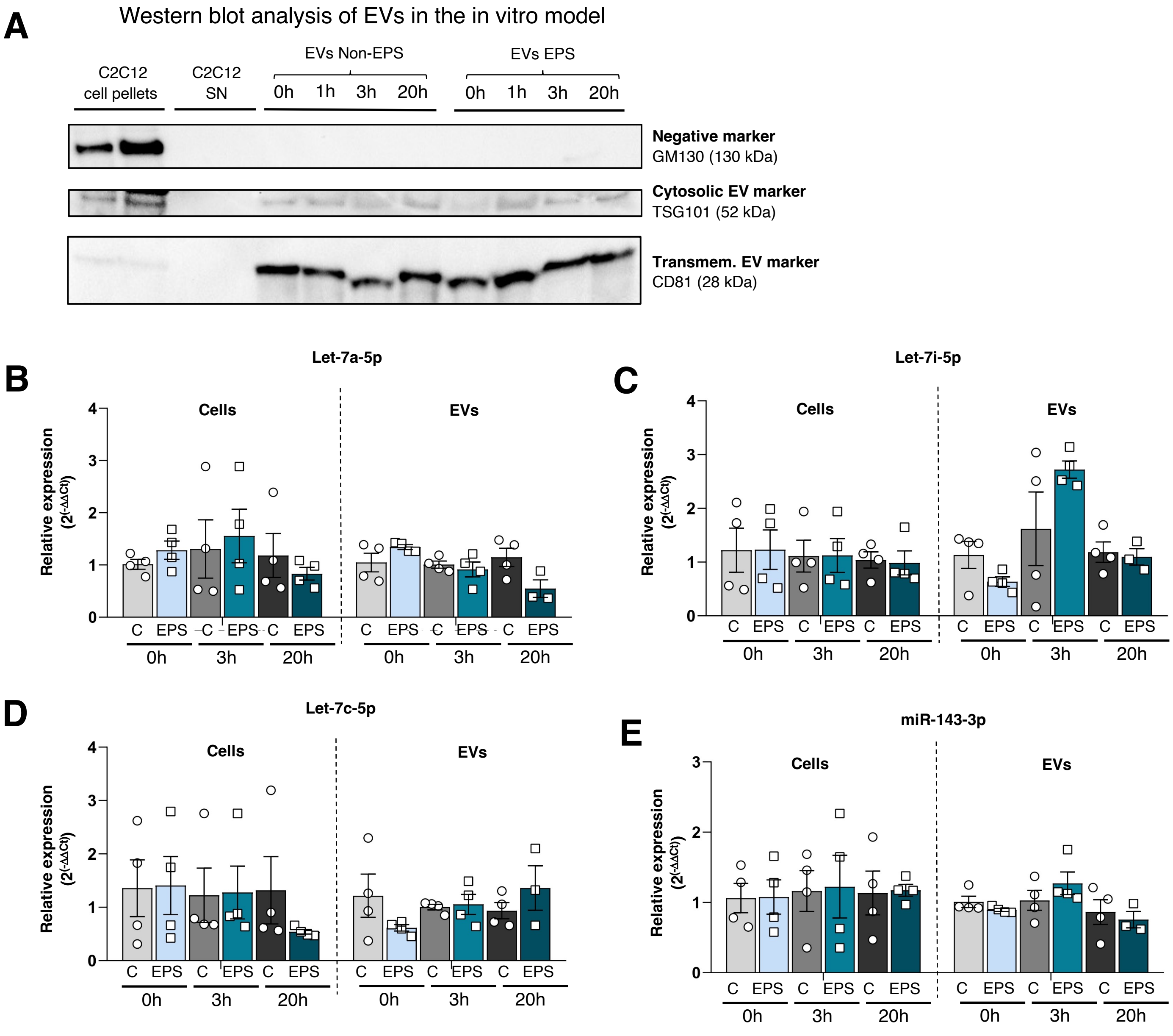


**Supplementary figure S4. Training-associated miRNA expression in C2C12 cells and EVs after EPS. (A)** Western blot analysis of the cytosolic EV marker TSG101, the transmembrane EV marker CD81 and the negative marker GM130 in EVs purified from culture media (n = 4), corresponding cell lysates (C2C12 cell pellets, n = 2) and culture media without EVs (C2C12 SN, n =2). EVs of non-EPS and EPS cells were compared at 0, 1, 3 and 20 hours. **(B-E)** Relative expression levels (2^(-ΔΔCt), measured by qPCR, of let-7a-5p, let-7i-5p, let-7c-5p and miR-143-3p in C2C12 cells and EVs, at 0, 3 and 20 h after the EPS in non-EPS (C) and EPS cells (EPS). Data are represented as mean ± SEM. Each dot represents a cell replicate (1 well) (n = 4/group). The statistical significance of 2^(-ΔΔCt) miRNA expression was calculated using the T-Student test for independent samples.

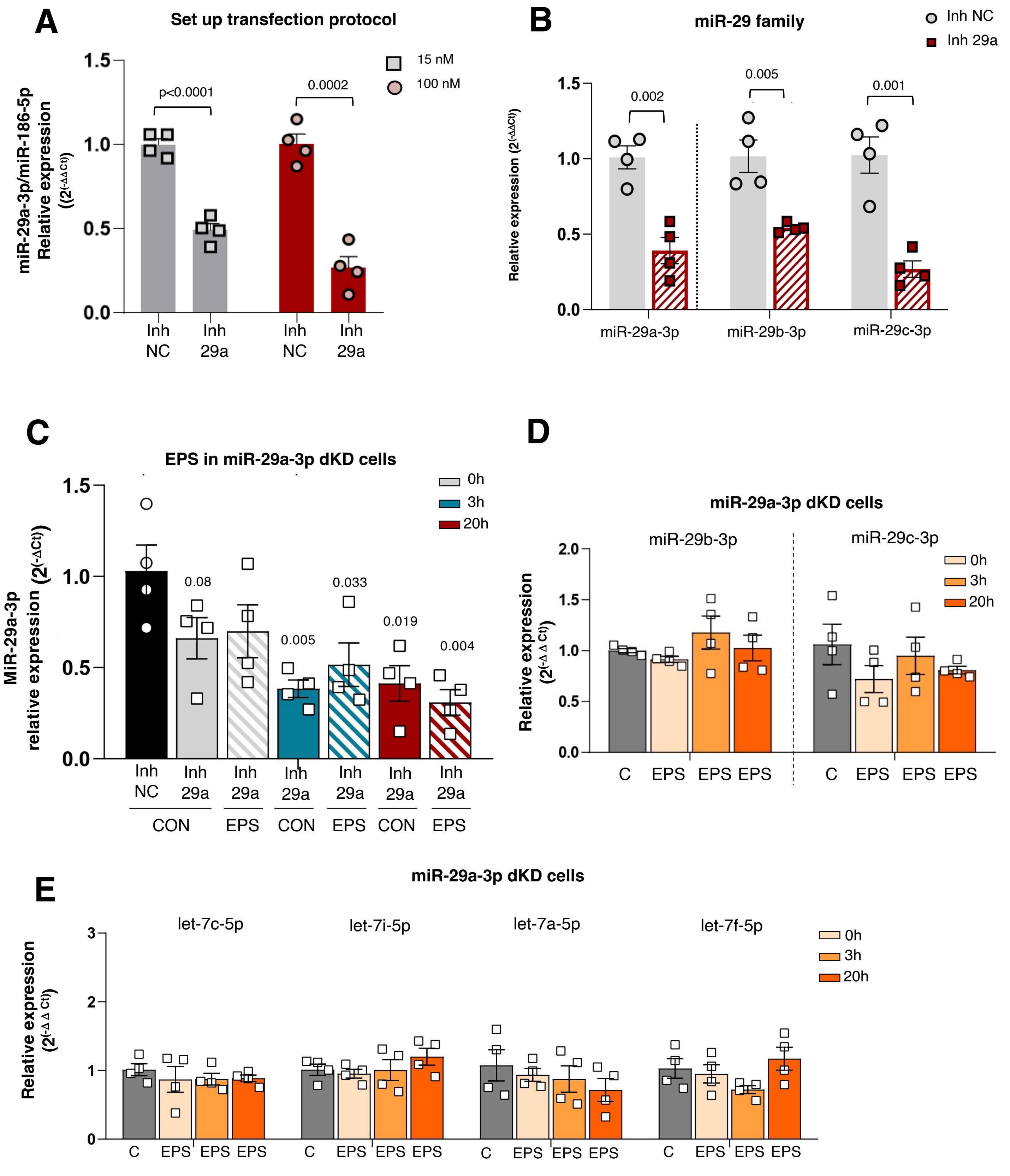


**Supplementary figure S5. Partial inhibition of miR-29a-3p and effect of EPS in C2C12 miR-29a-3p knockdown (dDK) cells. (A)** Efficacy of miR-29a-3p inhibition in C2C12 cells. Two different concentrations of the inhibitor and of the negative control (NC) were tested: 15 and 100 nM. The expression levels of miR-29a-3p (2^(-ΔΔCt)) were measured by qPCR and compared with the expression of the endogenous miRNA miR-186-5p. **(B)** Effect of electrostimulation in miR-29a-3p expression in dKD cells 0, 3 and 20h after EPS. The expression levels of miR-29a-3p (2^(-ΔCt)) were measured by qPCR. **(C-D)** Relative expression levels (2^(-ΔΔCt)) of miR-29b-3p, miR-29c-3p and the four let-7 family members in dKD cells 0, 3 and 20h after EPS in unstimulated cells (C) and stimulated cells (EPS). Expression levels were measured by qPCR and compared with the mean expression of all miRNAs. Data are represented as mean ± SEM. Each point represents one cell replicate (1 well) (n = 4/group). Statistical significance was calculated using the Student's t-test for independent samples. In figure B the statistical comparison was made with the first group (Inh NC CON group).

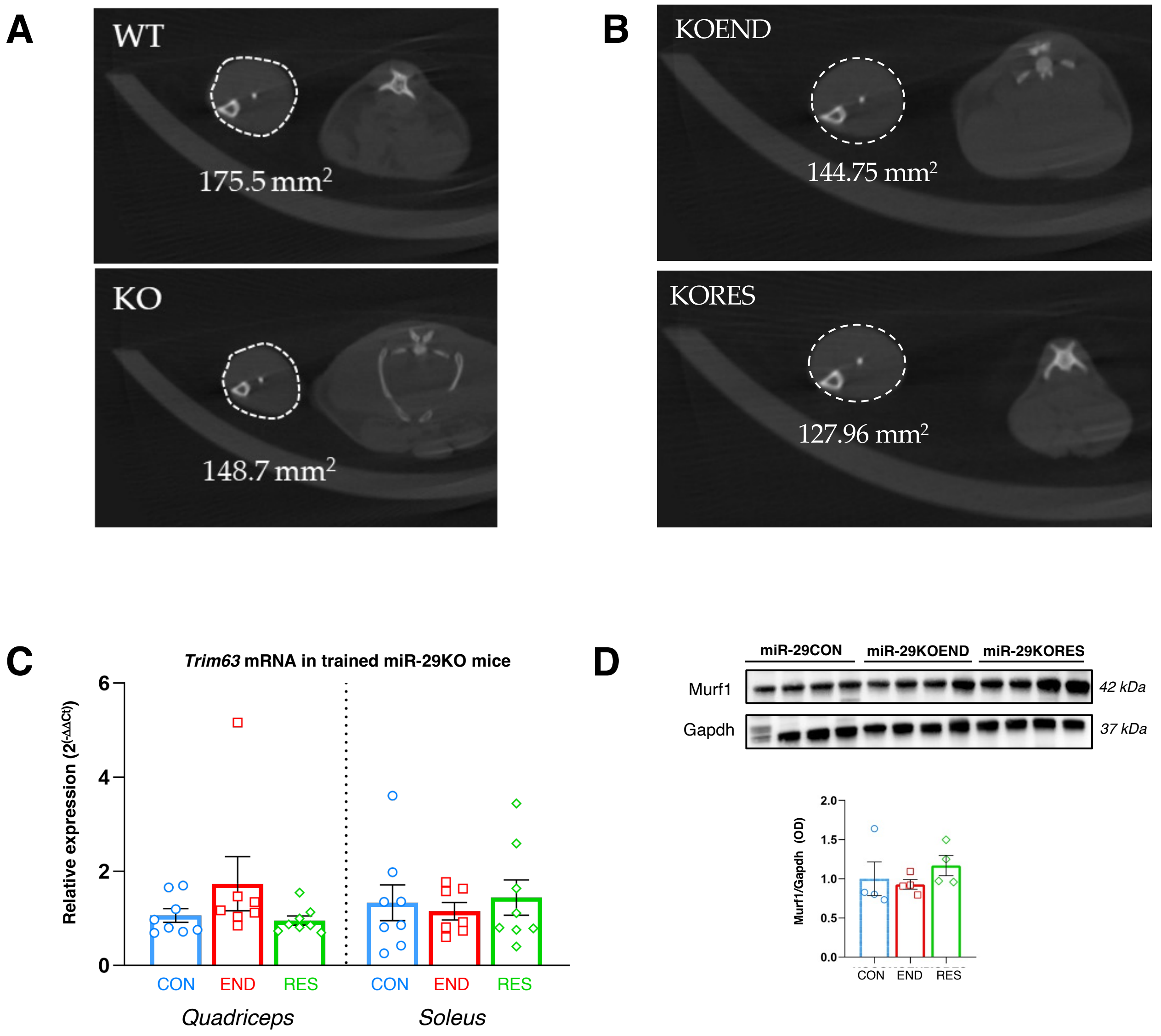


**Supplementary figure S6.** Analysis of muscle atrophy measured both macroscopically and molecularly in the quadriceps of miR-29KO trained mice at 16 weeks. **(A)** Micro-computed (CT) image of the right hind paw showing the total muscle area (mm2) of 1 WT and 1 miR-29KOCON. **(B)** Micro-computed (CT) image of the right hind paw showing the total muscle area (mm2) of 1 miR-29KOEND and 1 miR-29KORES mouse. **(C)** Relative expression levels (2(-ΔΔCt)) of Trim63, measured by qPCR, and normalised using Gapdh as reference gene (KOCON, n = 8; KOEND, n = 7 and KORES, n = 8). **(D)** Representatives immunoblot of Murf1 and Gapdh proteins level in quadriceps of miR-29KO trained mice and quantification of Murf1/Gapdh levels (arbitrary units [OD]) in a histogram (KOCON, n = 4; KOEND, n = 4 and KORES, n = 4) Data are represented as mean ± SEM. Statistical significance was calculated using a one-way ANOVA test followed by a Tukey post hoc test

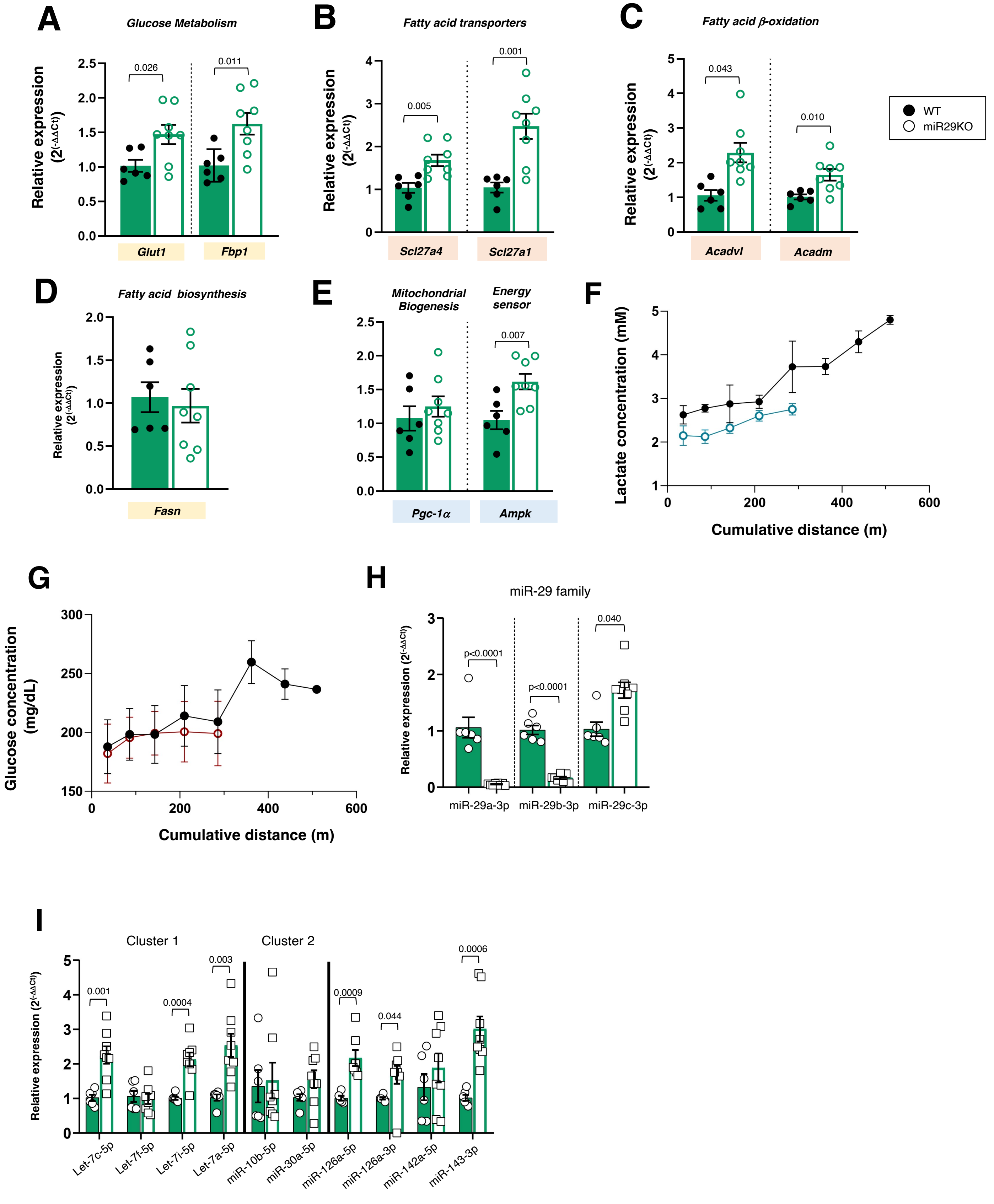


**Supplementary figure S7. Relative mRNA expressions of genes related to glucose metabolism (Glut1, Glut4, Pfkm and Fbp1), to fatty acid metabolism (Ffar4, Acadm, Acadvl, Slc27a1 and Slc27a4), energy state (Pgc-1α and Ampk) and anabolism (Mtor) in the quadriceps, soleus and liver of WT trained mice at 16 weeks.** Relative mRNA expression of each gene (2^(-ΔΔCt)) was normalized to *Gapdh* in the quadriceps and liver, and to *β-actin* in the soleus. Data are represented as mean ± SEM. Each point represents one mouse (CON, n = 6; END, n = 12 and RES, n = 8). Statistical significance measured by one-way ANOVA followed by Tuckey's test

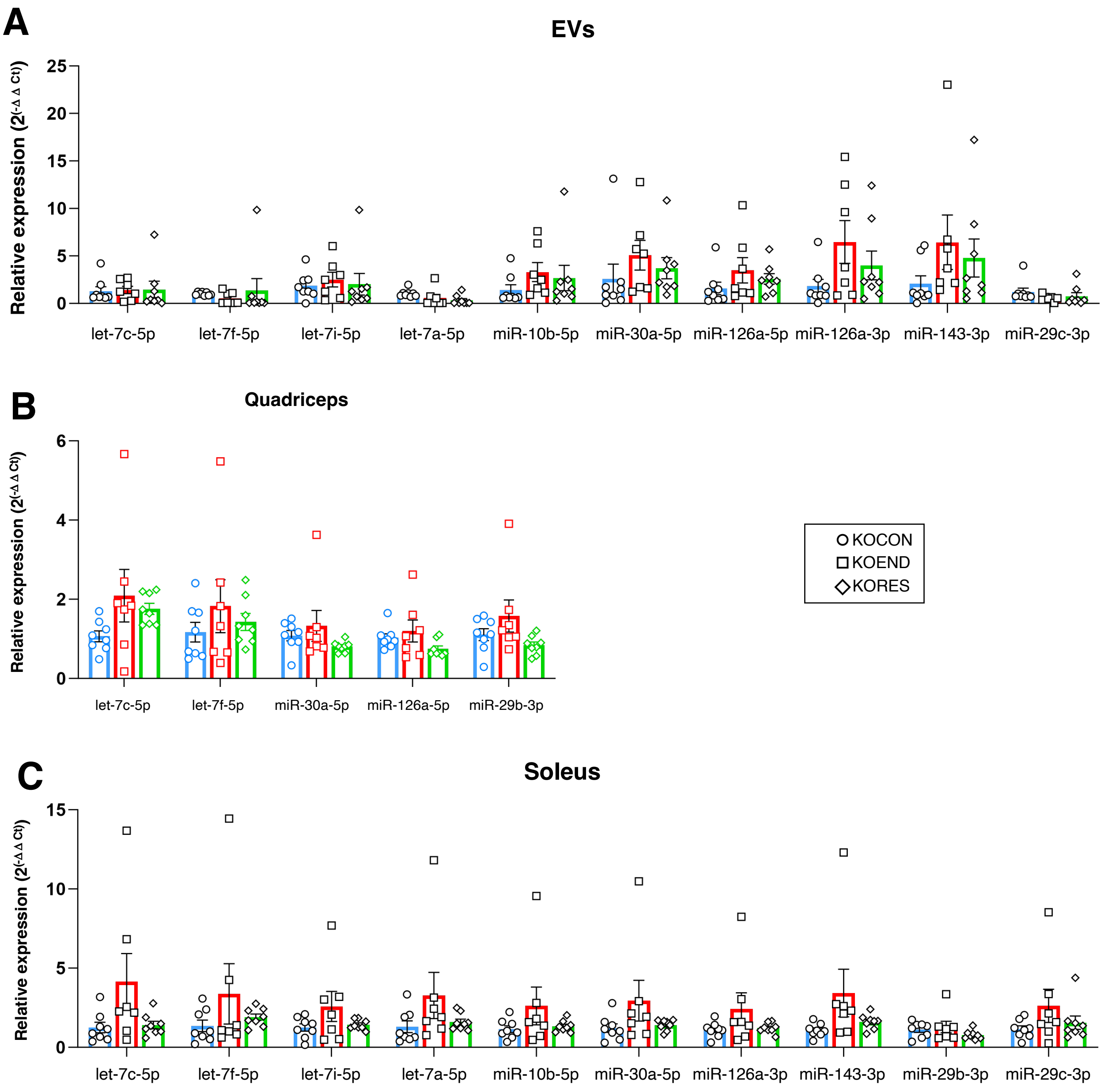


**Supplementary figure S8. Relative expression of training-associated miRNAs in miR-29KO trained mice. (A)** EVs. **(B)** Quadriceps. **(C)** Soleus. Relative expression levels (2^(-ΔΔCt)) were measured by qPCR and compared to Unisp6. Data are represented as mean ± SEM. Each point represents one mouse (CON, n = 8; END, n = 7; RES n = 8). Statistical significance measured by one-way ANOVA followed by Tuckey's post hoc test .

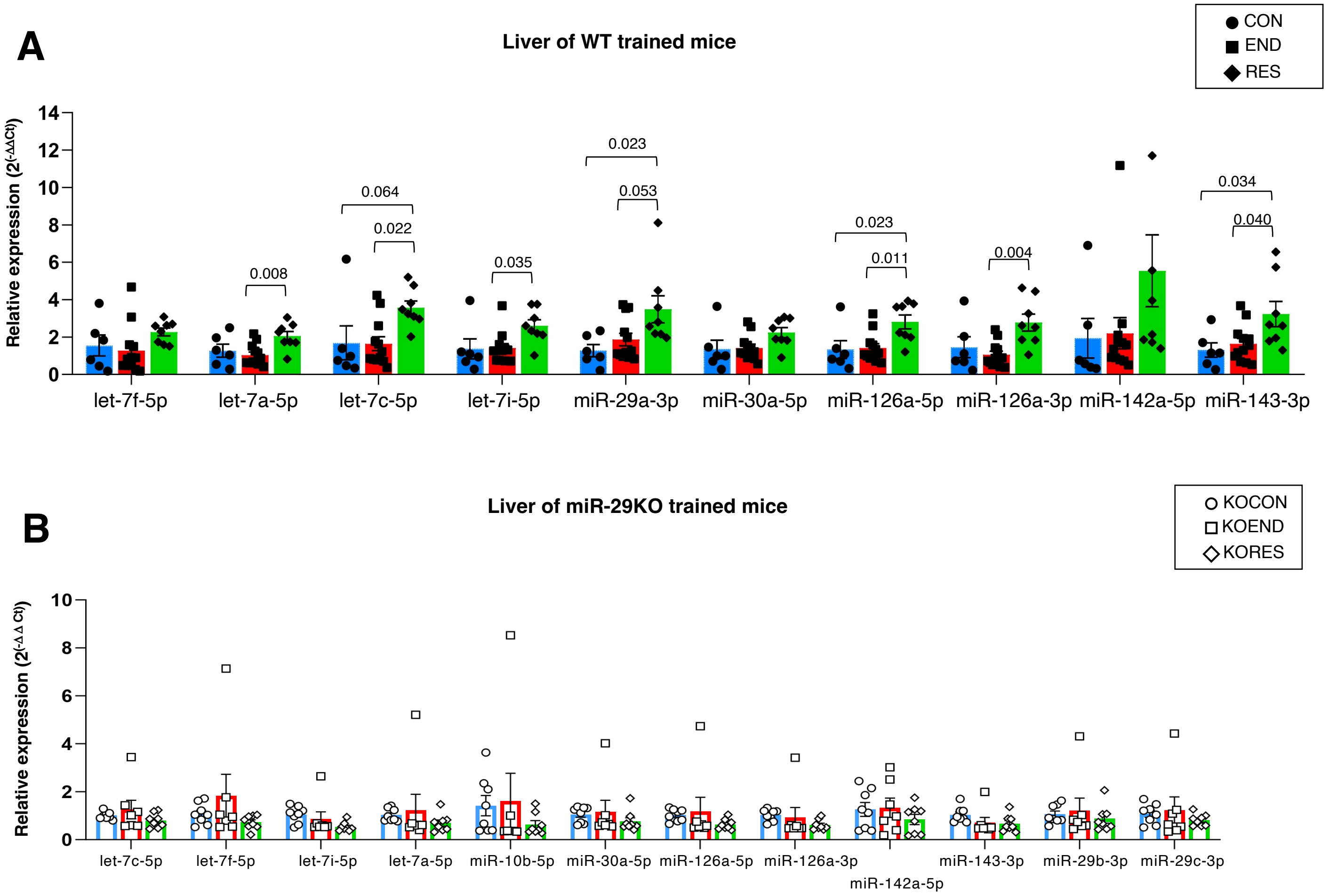


**Supplementary figure S9. Relative expression of training-associated miRNAs in liver of WT and miR-29KO trained mice.** **(A)** WT trained mice (CON, n = 6; END, n = 12; RES n = 8). **(B)** MiR-29KO trained mice (KOCON, n = 8; KOEND, n = 7; KORES n = 8). Relative expression levels (2^(-ΔΔCt)) were measured by qPCR and compared to Unisp6. Data are represented as mean ± SEM. Each point represents one mouse. Statistical significance measured by one-way ANOVA followed by Tuckey's post hoc test .

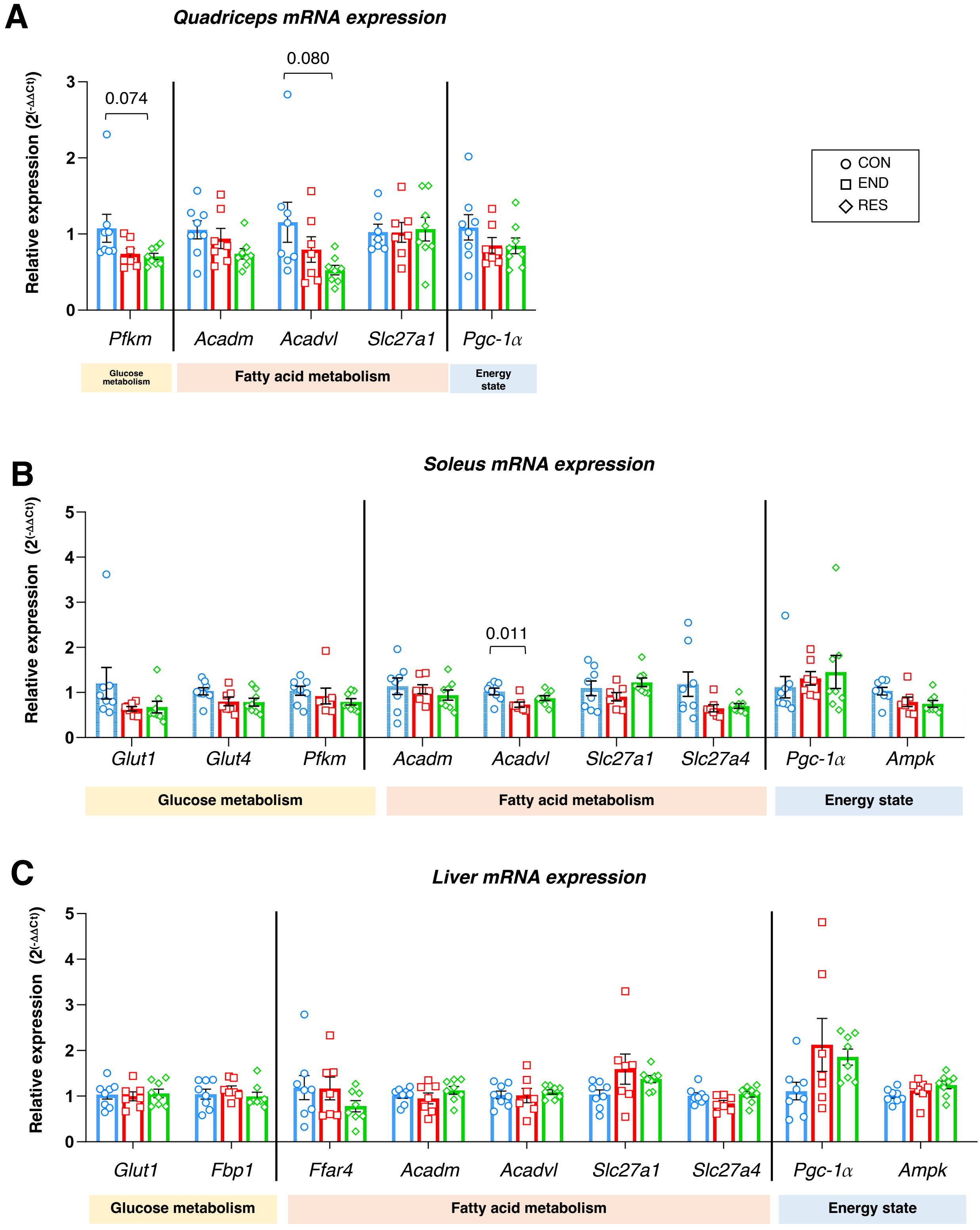


**Supplementary figure S10. Relative mRNA expressions of genes related to glucose metabolism (Glut1, Glut4, Pfkm and Fbp1), to fatty acid metabolism (Ffar4, Acadm, Acadvl, Slc27a1 and Slc27a4) and to energy state (Pgc-1α and Ampk) in the quadriceps, soleus and liver of miR-29KO trained mice.** Relative mRNA expression of each gene (2^(-ΔΔCt)) was normalized to *Gapdh* in the quadriceps and liver, and to *β-actin* in the soleus.. Data are represented as mean ± SEM. Each point represents one mouse (CON, n = 8; END, n = 7 and RES, n = 8). Statistical significance measured by one-way ANOVA followed by Tuckey's test

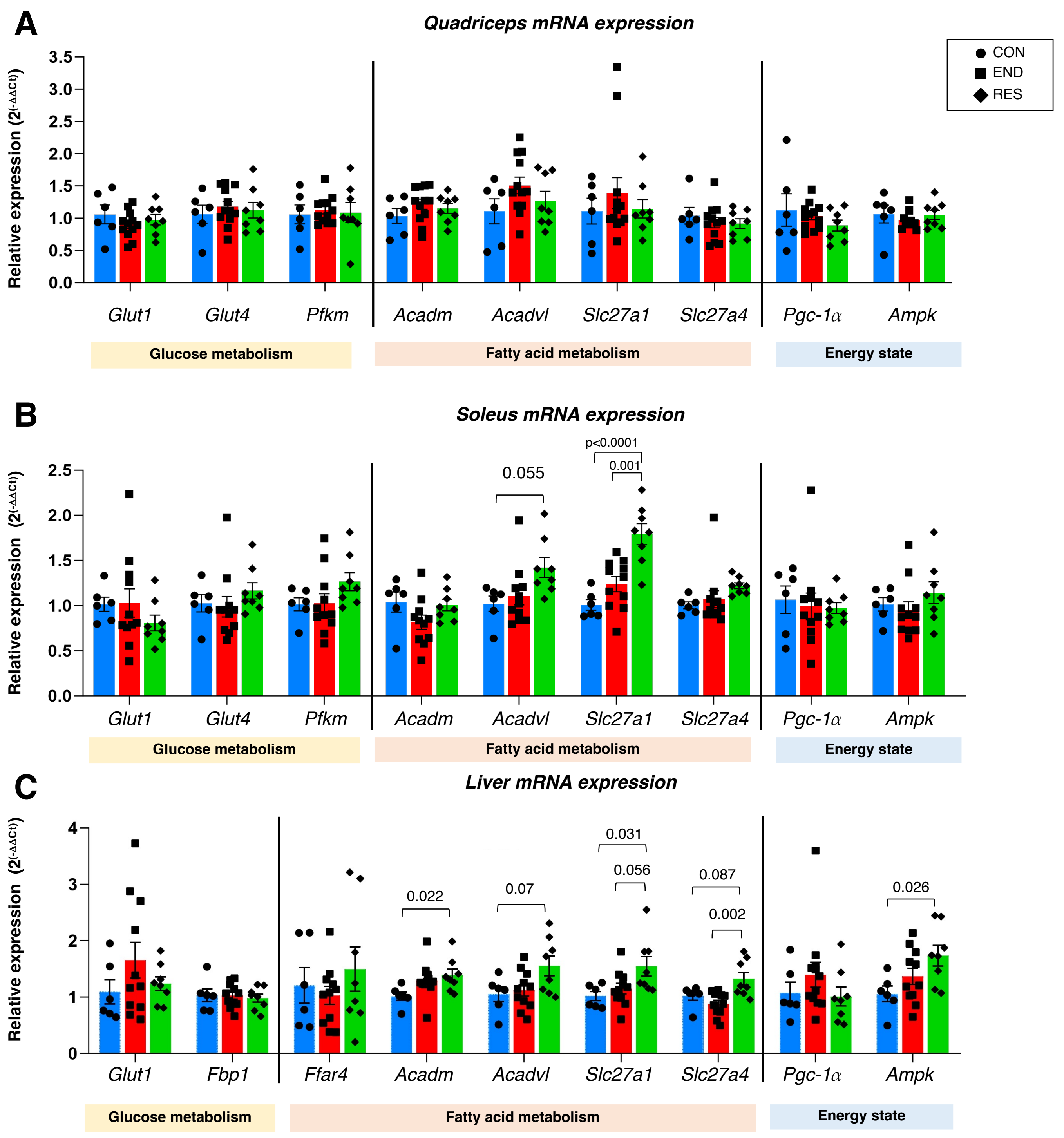


**Supplementary figure S11. Relative mRNA expressions of genes related to glucose metabolism (Glut1, Glut4, Pfkm and Fbp1), to fatty acid metabolism (Ffar4, Acadm, Acadvl, Slc27a1 and Slc27a4) and to energy state (Pgc-1α and Ampkin the quadriceps, soleus and liver of WT trained mice.** Relative mRNA expression of each gene (2^(-ΔΔCt)) was normalized to *Gapdh* in the quadriceps and liver, and to *β-actin* in the soleus. Data are represented as mean ± SEM. Each point represents one mouse (CON, n = 6; END, n = 12 and RES, n = 8). Statistical significance measured by one-way ANOVA followed by Tuckey's test

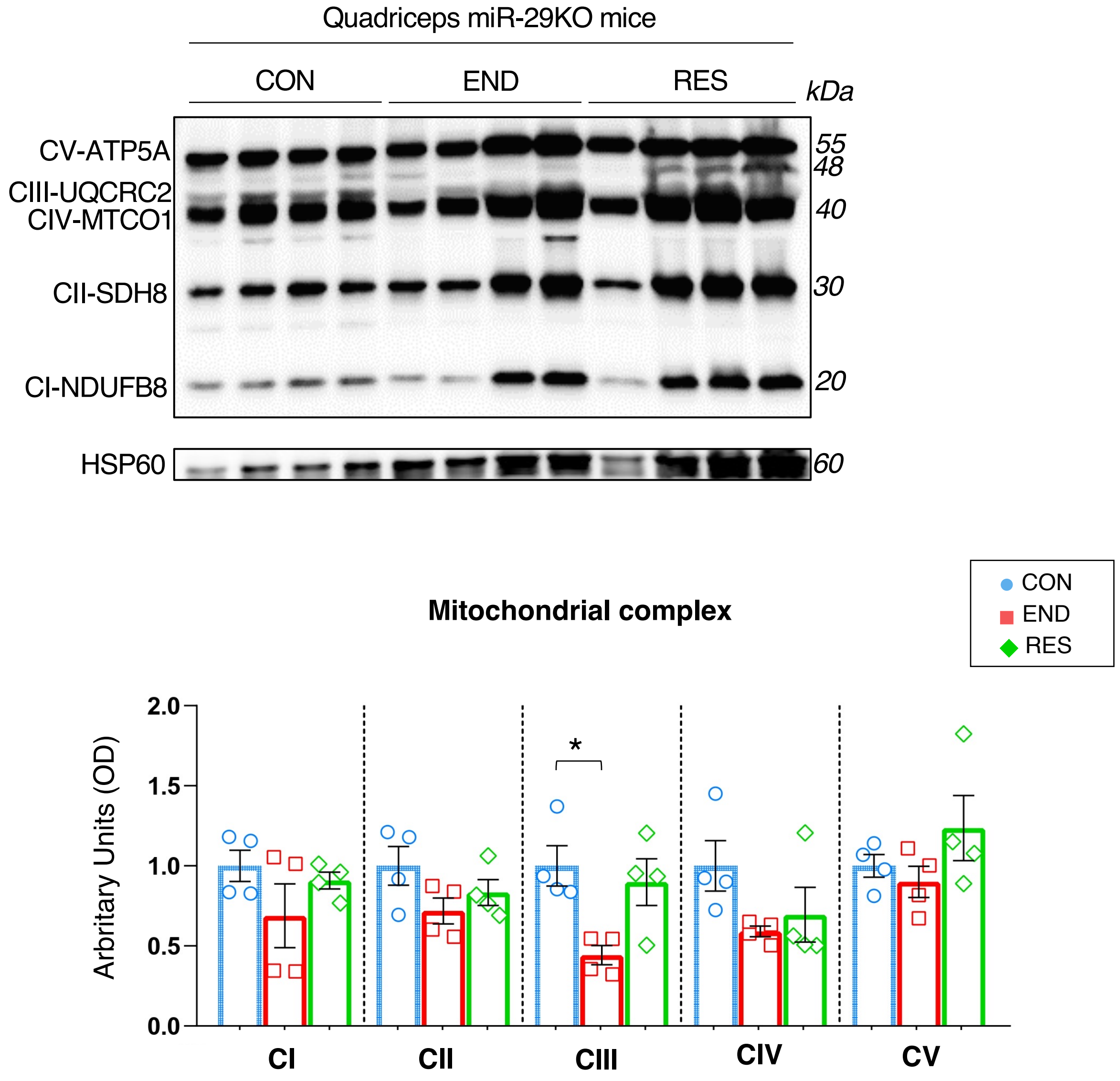


**Supplementary figure S12. OXPHOS protein levels in the quadriceps of miR-29KO trained mice at 16 weeks.** (A) Representatives immunoblot of OXPHOS and Hsp60 protein levels in quadriceps. (B) Quantification (arbitrary units [OD]) of the levels of each of subunits normalized to Hsp60 protein levels respectively in quadriceps of miR-29KOCON (n = 4), miR-29KOEND (n = 4) and miR-29KORES (n = 4). Data are represented as mean ± SEM. Each point represents one mouse. Statistical significance was calculated using a one-way ANOVA test followed by a Tukey post hoc test.
